# Generalizing predictions to unseen sequencing profiles via deep generative models

**DOI:** 10.1101/2021.05.06.443027

**Authors:** Min Oh, Liqing Zhang

## Abstract

Predictive models trained on sequencing profiles often fail to achieve expected performance when externally validated on unseen profiles. While many factors such as batch effects, small data sets, and technical errors contribute to the gap between source and unseen data distributions, it is a challenging problem to generalize the predictive models across studies without any prior knowledge of the unseen data distribution. Here, this study proposes DeepBioGen, a sequencing profile augmentation procedure that characterizes visual patterns of sequencing profiles, generates realistic profiles based on a deep generative model capturing the patterns, and generalizes the subsequent classifiers. DeepBioGen outperforms other methods in terms of enhancing the generalizability of the prediction models on unseen data. The generalized classifiers surpass the state-of-the-art method, evaluated on RNA sequencing tumor expression profiles for anti-PD1 therapy response prediction and WGS human gut microbiome profiles for type 2 diabetes diagnosis.

## Introduction

Predictive models relying on genomic signatures and biomarkers often suffer significantly inferior performance in the independent validation on external data sets in biomedical research such as disease diagnostics, prognostics, drug discovery, and precision medicine, resulting in a contribution to reproducibility crisis^1–4^. Irreproducible models can lead to not only invalid conclusions misleading subsequent studies but also a substantial waste of time and effort for researchers trying to commercialize the models to benefit patients^5^. A major factor behind these failures is the lack of generalizability across studies, in each of which the number of the heterogeneous data points is insufficient to obtain statistical power to overcome the generalization barrier. In addition to the sample size, usually, there is a significant gap between source data that are used to train classifiers and target data that are used to evaluate the classifiers. One possible cause of the gap is the batch effect such as different sample cohorts, different lab environments, and differences in experimental protocols across studies^4,6^, which violates the assumption that source and target data are drawn from the same distribution.

In many real-world applications, trained systems fail to produce accurate predictions for unseen data with the shifted distribution. For example, illumination or viewpoint changes in data acquisition for an object detection system and noisier environments for a speech-to-text translation system could easily disrupt the desired outcome. To address this issue, domain adaptation algorithms have been proposed to better align source and target data in a domain-invariant feature space when knowledge of target domains is available during the training phase^7–9^. However, in practice, it is common that no clue on the target domain is provided. As a more ambitious goal, domain generalization studies focus on training a model generalizing to the unseen domain without any foreknowledge of the unseen domain. Recent studies proposed different ways of domain generalization such as extracting domain-invariant features^10–12^, leveraging self-supervised tasks to guide and learn robust representation^13^, simulating domain shift in meta-learning^14^, and adding perturbed samples^15,16^. Although these methods achieved promising performance on benchmark data sets, their requirements, such as having datasets from multiple source domains or sufficient enough for splitting and simulating domain shift, are often not satisfied in biomedical research where only a limited number of heterogeneous data points in a single source domain is available. Data augmentation techniques in the computer vision field show promising potential in improving classifiers by reducing overfitting to source data^17–19^. Especially, recent advances in deep generative models such as generative adversarial networks (GAN)^20^ allow generating visual contents that are indistinguishable from real ones and also augmenting image data to guide in finding better decision boundaries^17,19^. More recently, generative models have been utilized to augment medical images, including Magnetic Resonance Images (MRI)^21^, computed tomography (CT)^22^, and X-ray images^23^. However, there has been little effort in transferring the success in computer vision to biomedical sequencing data^24^, although some studies tried to leverage computer vision techniques by regarding non-image data as image data^25–29^. Furthermore, it is unclear whether augmentation of sequencing data could overcome the generalization barrier across different studies.

In this study, DeepBioGen, a data augmentation procedure that establishes visual patterns from sequencing profiles and generates new sequencing profiles capturing the visual patterns based on conditional Wasserstein GAN, is proposed to enhance the generalizability of the prediction models to unseen data. DeepBioGen outperforms other augmentation methods in generalizing classifiers to unseen data. Also, the classifiers generalized by DeepBioGen surpass state-of-the-art classifiers that are designed to work on unseen profiles when tested on two scenarios: devising a prediction model for immune checkpoint blockade (anti-PD1) responsiveness in melanoma patients based on RNA sequencing (RNA-seq) data and building a diagnostic model for type 2 diabetes based on whole-genome metagenomic sequencing data. DeepBioGen source code is free and available at https://github.com/minoh0201/DeepBioGen.

## Results

### Formation and augmentation of visual patterns of sequencing profiles

Sequencing profiles, such as RNA-seq measurements of gene expression levels, consist of numerical values that indicate the activity of thousands of genes in different samples or patients. While many statistical methods such as multivariate linear regression assume that variables are independent of one another, in reality, genes’ activities are highly correlated^30^. In DeepBioGen, to take into account and visually formalize the interactivity of related genes, similar features in the profiles were clustered together, presenting visible patterns after converting numerical values to colors (Figure 1a; See Methods section). Subsequently, a conditional Wasserstein GAN equipped with convolutional layers to capture the local visual patterns was implemented to augment the sequencing profiles conditioned on class labels. During the augmentation phase, multiple GANs were initialized and trained with different random seeds to promote diversity in the augmented data points (Figure 1b).

**Figure 1.**
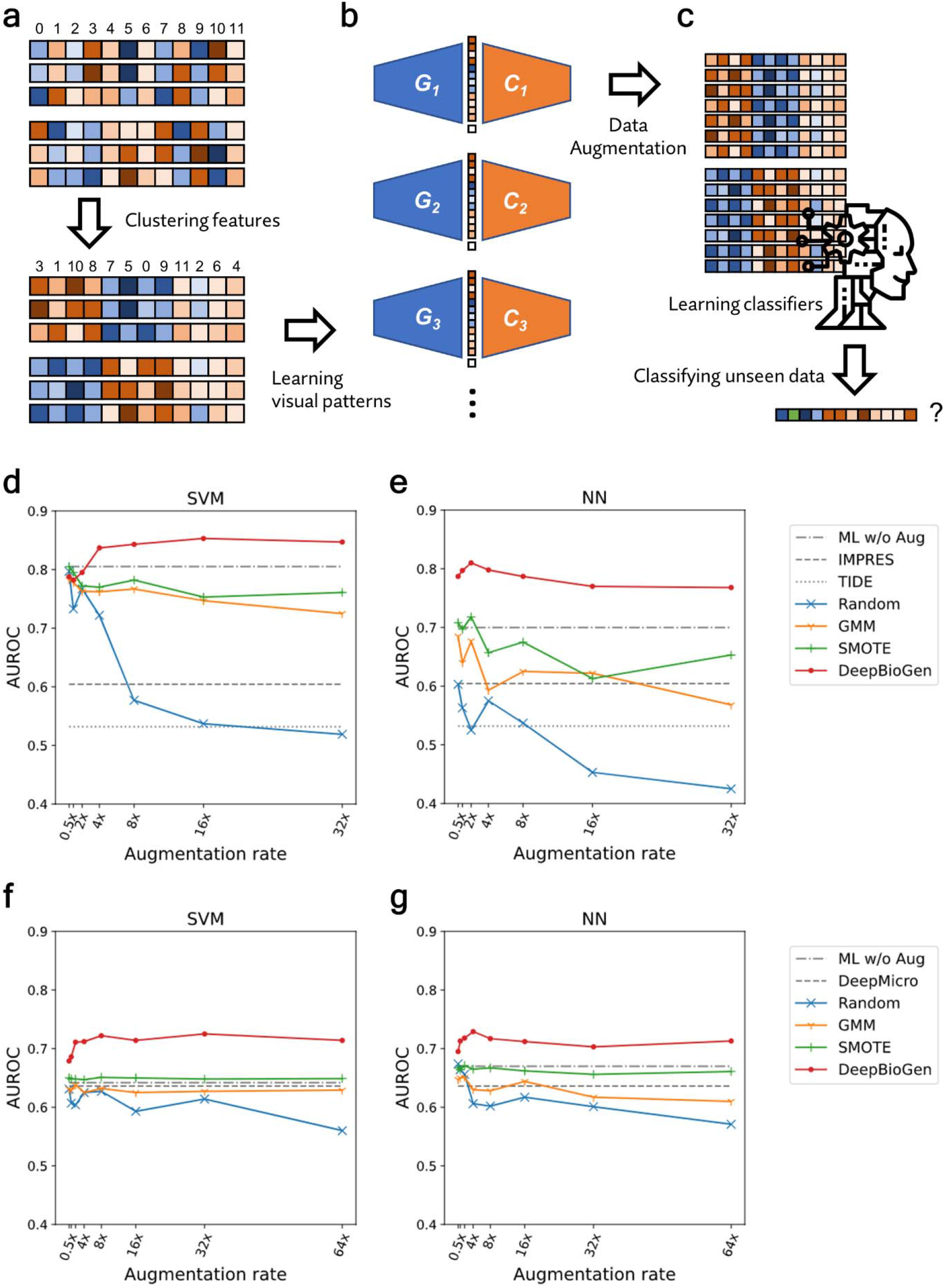
DeepBioGen, a sequencing profile augmentation procedure that generalizes classifiers to enhance prediction performance on unseen data. **a**, Feature-wise clustering of sequencing profiles to form perceptible visual patterns. **b**, Training multiple conditional Wasserstein GANs equipped with up-convolutional and convolutional layers. **c**, Generating augmented data from the multiple generators of GAN models and learning classifiers based on the augmented data along with source data to predict unseen data. **d-e**, Results of anti-PD1 therapy response prediction on unseen data by the state-or-the-art and baseline classifiers (gray) and by classifiers generalized with DeepBioGen (red), SMOTE (green), GMM (yellow), and Random augmentation (blue); Classification algorithms: Support Vector Machine (SVM) and Neural network (NN) which is a multi-layer perceptron; Evaluation metric: Area under the receiver operating characteristics (AUROC). **f-g**, Results of type 2 diabetes prediction on unseen data.

To inspect the visual quality of augmented data, two different sequencing profiles were used to train the generative models: one is RNA-seq expression profiles of melanoma patients, and the other is gut microbiome profiles of type 2 diabetes patients (See ‘sequencing profiles and pre-processing’ in Methods section and Supplementary Table S1 for details). Visual assessment showed that the augmented profiles preserved the boundaries of the clustered features and within-cluster color patterns in the same manner as source data. It is also difficult to distinguish an augmented profile from source data without the original tag (Supplementary Figure S1 and S2).

### Generalized classification on unseen sequencing profiles

The augmented data derived from the multiple generators of GANs was injected into training data along with the source data. The training data was used to train three machine learning classifiers, support vector machine (SVM), an artificial neural network (NN), and random forest (RF) (Figure 1c). The classifiers were trained to predict non-responders of cancer immunotherapy (anti-PD1) based on RNA-seq gene expression profiles or type 2 diabetes based on human gut microbiome profile (See method section and Supplementary Table S1).

To validate the generalizability of the classifiers, test (unseen) data were secured from studies that are independent of the source studies. Classification performances on test data were evaluated using an area under the receiver operating characteristics (AUROC) and an area under the precision-recall curve (AUPRC). State-of-the-art predictors, TIDE^31^ and IMPRES^32^ for predicting patient response to anti-PD1 therapy, and DeepMicro^33^ for using deep representations of microbiome data to predict disease states, were compared to DeepBioGen. Besides, widely-used data augmentation techniques, such as Gaussian Mixture Model (GMM)^34^ and Synthetic Minority Over-sampling Technique (SMOTE)^35^, were used to generate augmented data for comparison. The classifiers trained only on source data were used as the baseline comparison.

Remarkably, DeepBioGen-based classifiers surpass not only state-of-the-art classifiers but also classifiers that are trained on augmented data generated by different augmentation methods in both immunotherapy response (Figure 1d-e, Supplementary Figure S3, and Supplementary Table S4) and diabetes predictions (Figure 1f-g, Supplementary Figure S3, and Supplementary Table S5). Notably, even though DeepBioGen-based classifiers have no clue of test data, it outperforms Gide et al.’s immune marker classifier (AUROC=0.77) that directly leverages the test data through differential expression analysis^36^. Especially, DeepBioGen provides a stable performance boost to SVM and NN classifiers for both problems as the augmentation rate increases. RF classifiers partially benefit from DeepBioGen, showing generally worse performance than SVM and NN classifiers (Supplementary Figure S4). Consistently, DeepBioGen reduces 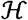-divergence between the source data and the test data more than other augmentation methods, which explains its better generalizability over others (Table 1).

**Table 1.**
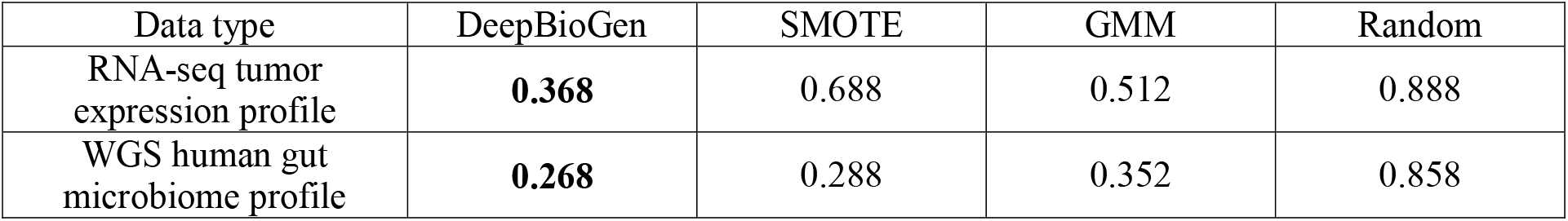
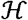-divergence between source and test data

| Data type | DeepBioGen | SMOTE | GMM | Random |
| --- | --- | --- | --- | --- |
| RNA-seq tumor expression profile | **0.368** | 0.688 | 0.512 | 0.888 |
| WGS human gut microbiome profile | **0.268** | 0.288 | 0.352 | 0.858 |

### Impact of visualized clusters and multiple generators

DeepBioGen uses the elbow method^37^ to estimate the optimal number of clusters and GANs (See ‘Formation of visual patterns from sequencing profiles’ in Methods section). To assess the ability of the approach in inferring the ideal parameters based on source data only, DeepBioGen models with a varying number of clusters or GANs were used to generate the augmented data for training classifiers. The classification results of unseen data show that the elbow method elicits an optimal or nearly optimal number of clusters and GANs in both immunotherapy response and diabetes prediction problems (Supplementary Figure S5-S8).

Notably, the number of clusters has more impact on classification performance than the number of GANs, suggesting that how sequencing data are clustered and thus presented visually plays a major role in improving the generalizability of DeepBioGen (Supplementary Figure S5-S8). Results also show that diverse generators of multiple Wasserstein GANs are more effective in diversifying the augmented sequencing data than a single generator, thus leading to better generalizability (Supplementary Table S3).

### Augmentations beyond the boundary of source data

To visualize how DeepBioGen augmented data to generalize classifiers, the source, augmented and test data were embedded to 2-dimensional space with t-distributed stochastic neighbor embedding (t-SNE) algorithm^38^. We note that t-SNE embedding preserves pairwise Euclidean distance in the 256-D space in both melanoma patient profiles and microbiome profiles (Pearson correlation r = 0.881 and r = 0.807).

In melanoma patient profiles, the source and test data are placed distantly, while within-cluster data points with different anti-PD1 responses are located closely in both data clusters (Figure 2a). The data embeddings were plotted separately for two classes, and an empirical outer boundary of the source data based on the outermost data points heading toward the test data was drawn with a red dotted line (Figure 2c and 2e). Interestingly, DeepBioGen generated data points (Figure 2b) beyond the outer boundaries of the source data cluster (Figure 2d and 2f), whereas other augmentation methods rarely produced data points that cross the boundaries (Supplementary Figure S7-S9).

**Figure 2.**
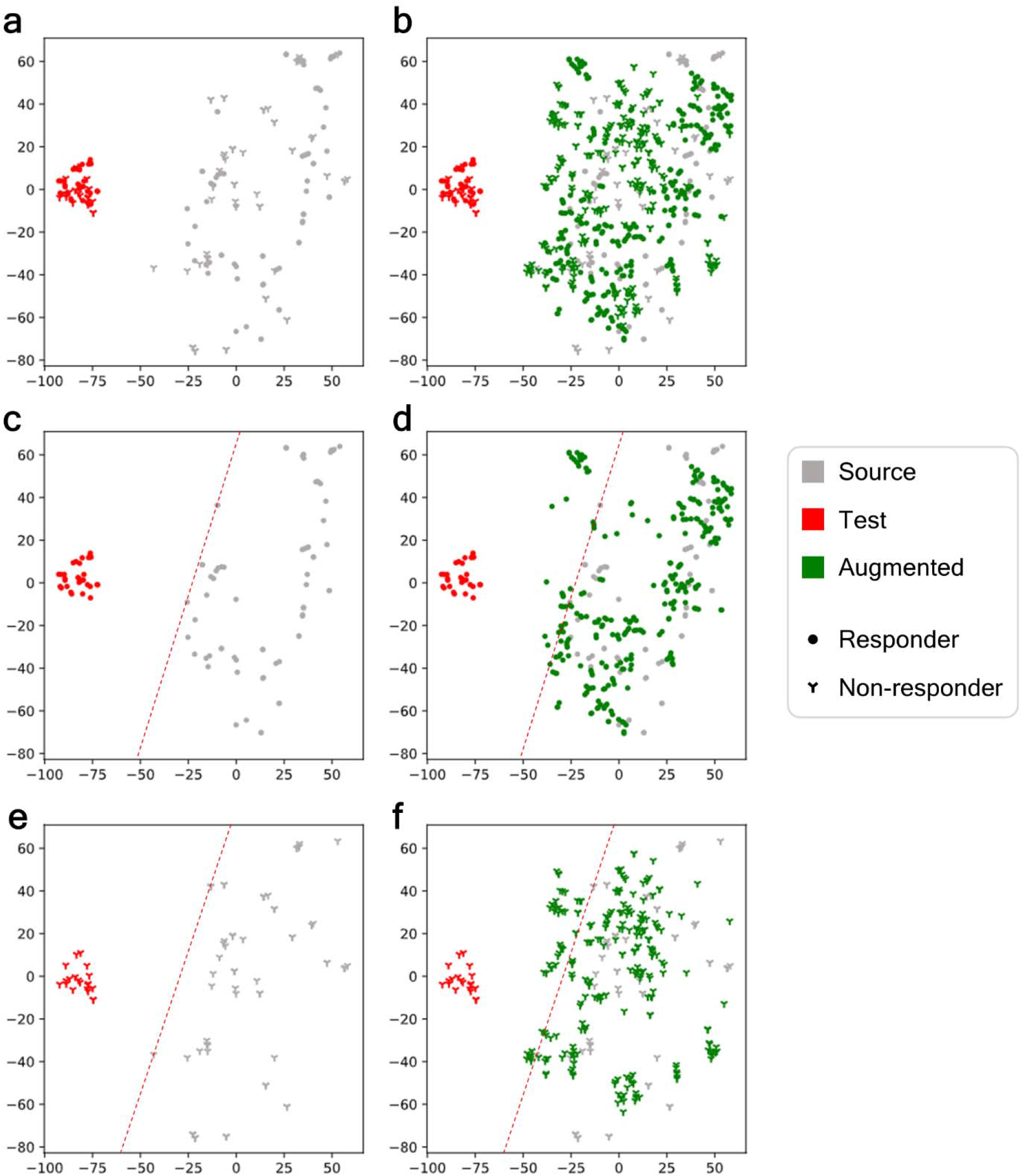
t-SNE visualization of augmented tumor expression profiles derived from DeepBioGen along with the source (grey), augmented (green), and test (unseen, red) data of melanoma patients treated with anti-PD1 therapy. **a**, The source and test data. **b**, The source, test, and augmented data. **c**, Responders of the source and test data; An empirical boundary of responders of source data (red dotted line). **d**, Responders of the source, test, and augmented data. **e**, Non-responders of the source and test data; An empirical boundary of non-responders of source data (red dotted line). **f**, Non-responders of the source, test, and augmented data.

In microbiome profiles of healthy controls and diabetic patients, the test data cluster resides in the side region of the source data cluster, thus depicting a moderately shifted distribution (Supplementary Figure S12). DeepBioGen produced augmented microbiome profiles across boundaries of the source data cluster. Particularly, the outermost augmented data points beyond the source boundaries are closely placed with test data points that cross the border (Supplementary Figure S12), while other methods rarely generate data points overpassing the boundaries (Supplementary Figure S13-S15).

### Progression-free survival analysis of predicted anti-PD1 treatment responders

For the predicted responder (PR) and non-responder (PNR) patients to anti-PD1 treatment determined by DeepBioGen-supported SVM classifier, progression-free survival analysis was conducted to estimate the clinical outcome. For comparison, state-of-the-art classifiers based on genomic signatures, IMPRES and TIDE, were evaluated with the same analysis. With the DeepBioGen classifier or IMPRES, the PR group has a significantly longer progression-free survival rate compared to the PNR group (Figure 3a and 3b), whereas the two TIDE predicted groups do not show a significant difference.

**Figure 3.**
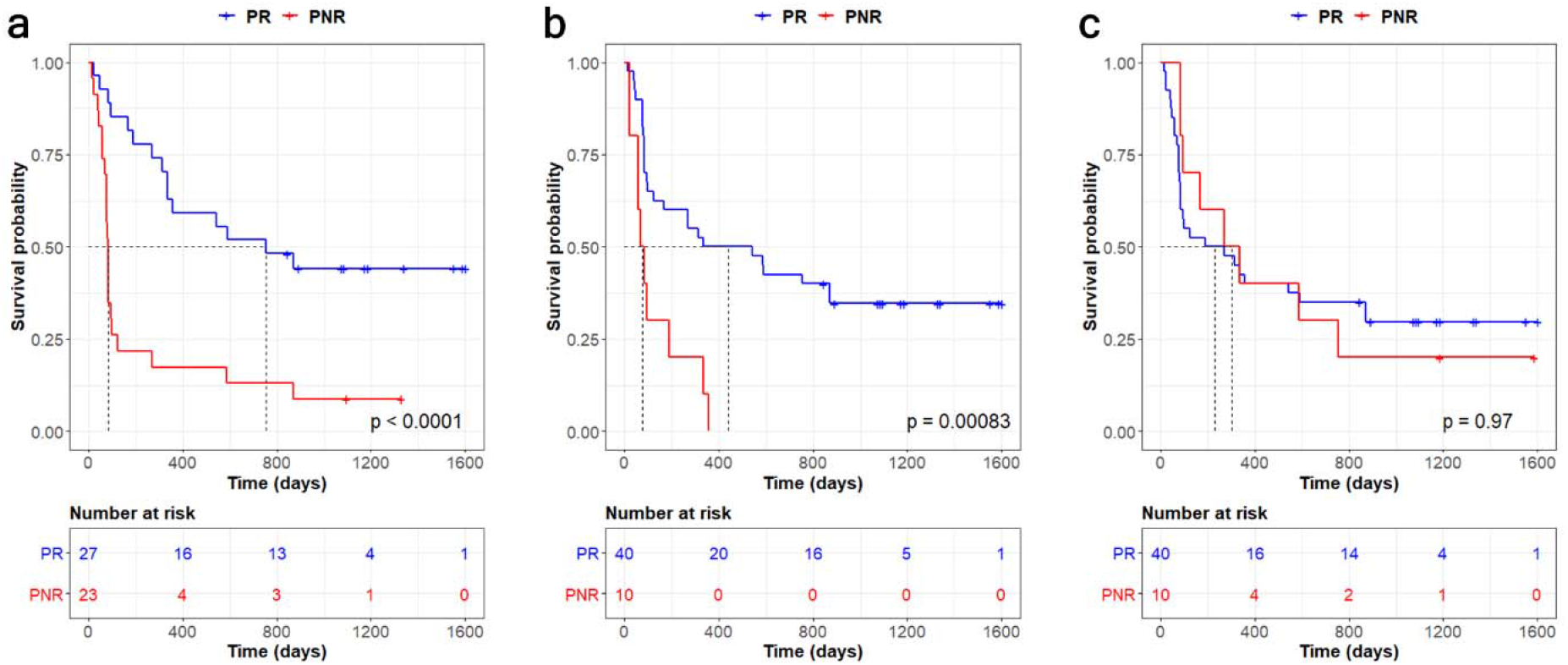
Kaplan-Meier plots of progression-free survival for predicted responder (PR) and nonresponder (PNR) patients determined by three classifiers. **a**, generalized SVM classifier with DeepBioGen augmentations. **b**, IMPRES. **c**, TIDE.

Importantly, the median survival time of PRs classified by the DeepBioGen classifier was 755 days (95% CI [335, N/A]), compared to 440 days (95% CI [125, N/A]) for the IMPRES classified PRs. Also, the DeepBioGen classifier tends to be more sensitive in predicting responders than IMPRES, likely posing a lower risk of unnecessary treatment suggestions often accompanied by unnecessary side effects (Figure 3 and Table 2).

**Table 2.** Summary statistics for progression-free survival analysis

| Classifier | Prediction | N | Median survival time (days) | 95% CI | MR<sup>*</sup> | HR<sup>**</sup> | 95% CI | P-value |
| --- | --- | --- | --- | --- | --- | --- | --- | --- |
| DeepBioGen-SVM | PR | 27 | 755 | [335, NA] | 9.21 | 3.72 | [1.88, 7.36] | < 0.001 |
|  | PNR | 23 | 82 | [76, 125] |  |  |  |  |
| IMPRES | PR | 40 | 440 | [125, NA] | 5.71 | 3.47 | [1.66, 7.49] | 0.002 |
|  | PNR | 10 | 77 | [58, NA] |  |  |  |  |
| TIDE | PR | 40 | 231 | [82, 870] | 0.76 | 0.99 | [0.45, 2.17] | > 0.9 |
|  | PNR | 10 | 303 | [96, NA] |  |  |  |  |

<sup>\*</sup>Median ratio; <sup>\*\*</sup>Hazard ratio

## Discussion

DeepBioGen provides a framework for effective data augmentation in sequencing profiles that can be used to boost the training data and improve the performance of prediction models on unseen data. It adversarially learns multiple generative models that capture visual signals from source data. With multiple generators, DeepBioGen generates realistic augmented data beyond the boundary of the source domain. The augmented data can be used to amplify training data and train classifiers resilient to unknown domain shifts. Consequently, DeepBioGen can improve the transferability and reproducibility of the prediction models without any knowledge of unseen data.

The stable performance over augmentation rate with DeepBioGen was observed with SVM and NN applications but not in RF. One of the possible reasons might be related to the fact that SVM and NN are the methods approximating decision boundary, while RF is aggregating results from bootstrapping. Augmentations could introduce relatively fewer variations on smoothing decision boundaries as source data always preserves its contribution to the decision boundaries, whereas augmentation together with bootstrapping could have a negative impact on performance because bootstrapped samples may have too little purity to reasonably optimize the best split.

DeepBioGen is unique as it takes input sequencing profiles in machine-understandable visual form, while visualization of sequencing data (e.g. heatmap of differentially expressed genes) has been typically used to present findings in a human-understandable manner. One potential advantage of feeding DeepBioGen with visually recognizable data is that visual patterns difficult to be identified with human eyes may be captured and characterized in embedding space.

Even with a limited amount of source data, DeepBioGen can alleviate batch effects of independent studies without details for batch correction such as sample cohorts, lab environments, and experimental protocol, by reducing the gap between the source and unseen data. Also, DeepBioGen is highly extensible to other biological data whose feature dependency is not negligible.

One limitation of this study is that there is no theoretical guarantee of improvement for other datasets except the ones tested in this paper. For example, if domain shift is severe enough so that there is no similarity between source and test data at all, the augmentations may not be helpful to improve performance. Also, depending on how hard the classification problem is and how stable the optimization of hyper-parameters in the experiment is, given the limited data, the performance on unseen data could be impacted. Conducting cross-validation for different splits within source data or combined source and test data could be a good way to fathom these properties. We conducted 5-fold cross-validation on source data and repeated ten times with different random splits to evaluate the prediction performance (Table S6 and S7). We also conducted 5-fold cross-validation on the combined source and test data with random splits for ten times (Table S8 and S9). We found that the performance of different classification models was significantly better than random guessing and the performance variations could be reduced by adding more data points.

It is also worth mentioning that there should be some sort of similarity between the source and unseen data for classifiers to take the advantage of the augmentations. Without similarity, it is hard to expect an improvement of performance on unseen data. We tried to find underlying common properties of source and unseen data to understand the reason that classifiers perform relatively well on unseen data with limited data points in high-dimensional space. We extracted the most important 25 features out of the 256 features from both the source and test data (based on Gini index with decision tree algorithm to predict class labels) and checked the number of overlapping features. Interestingly, we found statistical significance in the overlaps: 12 features are overlapping for the tumor expression data set (one-tailed fisher’s exact test p = 0.00001) and 6 features for microbiome data (one-tailed fisher’s exact test p = 0.0231). This reveals that there are some shared properties between the source and test data which may in turn contribute to the success of using augmented data for improving classification performance.

In the future study, it is envisioned that the process of forming visual patterns from sequencing profiles can be learned with cutting-edge machine learning models toward the better formation of machine-understandable patterns.

## Methods

### Sequencing profiles and pre-processing

Clinical genomic data containing RNA-seq tumor expression profiles of melanoma patients and their responsiveness to anti-PD1 therapy were secured from three independent studies^36,39,40^ (Supplementary Table S1). Fifty samples in the most recent study^31^ were used as test data and the others were used as source data. RNA-seq read counts were normalized to transcripts per million (TPM) and then log2-transformed. To focus on genes related to primary mechanisms of tumor immune evasion, recently identified T cell signature genes^31^, such as regulators of T cell dysfunction and suppressors of T cell infiltration into the tumor, were selected out of 18,570 common genes across the studies. In total, 702 genes were considered as features of initial inputs.

Human gut metagenomic sequencing reads of type 2 diabetic patients and healthy controls were acquired from two independent studies: one on the Chinese cohort^41^ and the other on the European women cohort^42^ (Supplementary Table S1). Using MetaPhlAn2^43^, strain-level marker profiles were extracted from the metagenomic samples. In total, the number of common strain-level markers that are considered as initial features was 74,240. The European samples in the more recent study were used as test data and Chinese samples as source data.

### Formation of visual patterns from sequencing profiles

Each measurement in source data was standardized by subtracting the mean and dividing by the standard deviation. The same standardization was applied to test data using the mean and standard deviation of source data. To meet the dimensional requirement of the pre-defined input layer, the extremely randomized trees^44^ feature selection algorithm was applied to the source data to select 256 features. The k-means clustering algorithm was used to cluster features, minimizing squared Euclidean distances between centroid and within-cluster features. Based on the elbow point where the within-cluster sum of squared errors (WSS) starts to decrease significantly, the optimal number of clusters was determined to be 4 for RNA-seq tumor expression profiles and 6 for human gut microbiome profiles (Supplementary Figure S16). The selected features were then sorted and rearranged by cluster labels so that similar features are placed nearby. The features of test data were also rearranged in the same order.

### Augmentation of sequencing profiles based on their visual patterns

DeepBioGen captures local visual patterns of sequencing profiles by training conditional Wasserstein GAN, whose generator and critic networks are composed of up-convolutional and convolutional layers, respectively. The generator tries to generate realistic images enough to fool the critic, whereas the critic tries to assign higher values for real images than for generated images. During training, the generator and the critic progressively become better at their jobs by competing against each other. This adversarial training can be conducted by optimizing a minimax objective. Wasserstein distance (or Earth Mover) formulated by Kantorovich-Rubinstein duality is used in the objective term for better reaching Nash equilibrium^45^. Also, the gradient penalty is applied to the objective function to enforce the Lipschitz constraint, alleviating potential instability in the critic^46^. Generator function *G* and critic function *C* are conditioned on the class label *y* and the final objective function of conditional Wasserstein GAN is as follows:

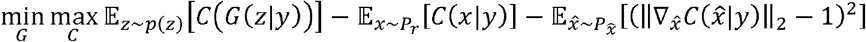

where *z* denotes a random noise vector derived from random noise distribution *p*(*z*), *x* a real profile derived from the real data distribution *P_r_*, and 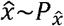 sampling uniformly along straight lines connecting the real data distribution *P_r_* and the output distribution of generator *P_g_* = *G*(*z*|*y*). The gradient penalty term directly constrains the norm of the critic’s output concerning its input, enforcing the Lipschitz constraint along the straight lines.

The architecture of neural networks that approximate generator function *G* and critic function *C* is illustrated in Supplementary Figure S17. The generator begins with two input layers, one for receiving a random noise vector and the other for a class label, followed by dense (64 units) and embedding layers (50 units). Embedded random noise vector and label vector are reshaped and concatenated. Subsequently, two up-convolutional blocks, composed of an up-convolutional layer, batch normalization layer, and Leaky ReLU activation layer, perform inverse convolution operations. Lastly, the final up-convolutional layer produces generated sequencing profile. Note that each sequencing profile is considered as a 1×256 pixel image in a single channel. Similarly, the critic has two input layers, one for sequencing profile and the other for a class label, which is embedded, reshaped, and concatenated onto the sequencing profile vector. The two consecutive convolutional blocks, each of which consists of a convolutional layer, Leaky ReLU activation, and dropout layer, are followed by the output layer with a single unit. Across the generator and critic, the alpha value of Leaky ReLU is set as 0.3, and the dropout rate is set at 0.3.

To achieve better generalization, multiple clones of the GAN are trained in the same way except for initial weights in the neural networks. The number of desired GANs is estimated by approximating modes of samples with the elbow method under the assumption that most modes are generated if the number of generators is at least as many as the number of modes in source data (Supplementary Figure S18). Individual generators produce the same number of augmented data points.

### Generalized predictions on unseen sequencing profiles

To generalize classifiers predicting clinical outcomes or disease states to unseen data, three classifiers, SVM, NN, and RF, were built on training data composed of source and augmented sequencing profiles. Hyper-parameters of the classifiers were optimized based only on source data with a 5-fold cross-validation scheme. Grid search was applied to explore hyper-parameter space (see details in Supplementary Table S2). With the best hyper-parameters, prediction models were trained on the pooled source and augmented data. The generalizability and performance of the prediction models were evaluated on the unseen test data using AUROC and AUPRC. The performance evaluation was repeated by gradually changing the augmentation rate indicating how many times the size of augmented data is of the source data.

For comparison, state-of-the-art classifiers designed to work on unseen data, including TIDE^31^, IMPRES^32^, and DeepMicro^33^, were evaluated on test data. TIDE predicts anti-PD1 responsiveness of melanoma patients based on genome-wide expression signatures of T cell dysfunction and exclusion. To satisfy its requirement, the test data without filtering out any genes from the original data was submitted to the TIDE response prediction web service. IMPRES is a predictor of anti-PD1 response in melanoma patients, which is a rule-based classifier manually built based on gene expression relationships between immune checkpoint gene pairs. Its source code was utilized to evaluate the performance of IMPRES on the test data. DeepMicro is a deep representation learning framework for improving predictors based on microbiome profiles. The source data was utilized to learn a low-dimensional representation of the microbiome data, and classifiers were then trained on the representation and evaluated on the test data. Furthermore, as an alternative to DeepBioGen, widely-used data augmentation approaches, including GMM^34^ and SMOTE^35^, as well as statistics-based random augmentation were evaluated. An independent GMM model was fitted for each class label, and the optimal number of components in the GMM model was estimated with the Bayesian information criterion (BIC). SMOTE derives the generated samples from linear combinations of nearest neighboring samples. Random augmentation draws data points from the normal distribution whose mean and standard deviation are the same as those of the source data, assigning an arbitrary class label. Also, as a baseline comparison, machine learning classifiers that are trained only on source data (i.e., no augmented data) were evaluated on test data.

To understand the impact of generalization on reducing the discrepancy between the source and test data, a classifier-induced divergence measure, 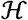-divergence, was determined with various classifiers. For a given set of binary hypotheses 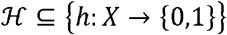, 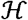-divergence is the largest possible difference between probabilities of being classified as 1 in source and test distributions^47,48^. More formally, the empirical 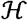-divergence can be written as:

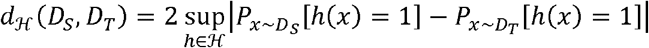

where *D_S_* and *D_T_* are the source and test data, respectively, and

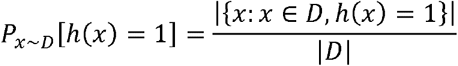

As a proxy of 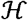 for each augmentation method, all classifiers trained on the augmented training data by varying an augmentation rate and classification algorithms were included in a set of binary hypotheses.

### Impact of multiple generators on the diversity of generated sequencing profiles

Wasserstein GAN may suffer less from mode collapse than infant GAN relying on Jensen-Shannon divergence in its loss term^45^. However, a single Wasserstein GAN may not be able to produce all modes of data, and it can be hypothesized that multiple Wasserstein GANs may increase the diversity of augmented sequencing profiles. To evaluate the diversity of the augmented profiles generated with multiple Wasserstein GANs, the adapted inception score is used. Originally, the inception score was introduced to evaluate the quality and diversity of generated images based on the predicted class probability distributions derived from a pre-trained Inception v3 model^49^. More recently, Gurumurthy et al. suggested a modified inception score considering within-class diversity of the generated data^50^, and this scoring method is used in the current evaluation. Also, according to the note that non-ImageNet data generator should not be evaluated by the Inception v3 classifier^51^, it is replaced with the best performing baseline-classifier trained only on source data. Consequently, the adapted inception score ranges from 1 to 2, and the higher the score, the better the diversity and quality of the augmented profiles.

### t-SNE visualization of the augmented data

To visualize how augmented data is arranged in a high-dimensional space, the augmented data along with source and test data was embedded into a 2-dimensional space using t-SNE. Also, a class-specific boundary of the source data cluster facing the test data cluster in the embedded space was drawn with one or two straight lines through the outermost data points of the source data cluster.

### Progression-free survival analysis

The Kaplan-Meier plots were drawn to conduct progression-free survival analysis for predicted responder and non-responder patients. For each classifier, a receiver operating characteristic (ROC) curve was used to determine the cut-off value of predictions. The closest point from (0, 1) on the ROC curve was chosen, at which the threshold well balancing true positive rate and false-positive rate is identified. The log-rank test was used to validate statistical significance.

