## Supplementary materials for "Generalizing predictions to unseen sequencing profiles via deep generative models"

Supplementary materials of
**Generalizing predictions to unseen sequencing profiles via deep generative modeling**

Min Oh^1^ and Liqing Zhang^1, *^

^1^Department of Computer Science, Virginia Tech, Blacksburg, VA, USA

**Supplementary figures:**

- **Figure S1.** Visualization of tumor gene expression profiles of melanoma patients.
- **Figure S2.** Visualization of microbiome marker profiles of diabetic patients and healthy controls.
- **Figure S3.** Prediction performance on unseen data (AUPRC).
- **Figure S4.** Prediction performance on unseen data with Random forest (RF) classifier.
- **Figure S5.** Anti-PD1 therapy response prediction performance (AUROC) on unseen data by varying the number of visual clusters and that of GAN models.
- **Figure S6.** Anti-PD1 therapy response prediction performance (AUPRC) on unseen data by varying the number of visual clusters and that of GAN models.
- **Figure S7.** Type 2 diabetes prediction performance (AUROC) on unseen data by varying the number of visual clusters and that of GAN models.
- **Figure S8.** Type 2 diabetes prediction performance (AUPRC) on unseen data by varying the number of visual clusters and that of GAN models.
- **Figure S9.** t-SNE visualization of augmented tumor expression profiles derived from Random augmentation along with the source and test (unseen) data of melanoma patients treated with anti-PD1 therapy.
- **Figure S10.** t-SNE visualization of augmented tumor expression profiles derived from GMM along with the source and test (unseen) data of melanoma patients treated with anti-PD1 therapy.
- **Figure S11.** t-SNE visualization of augmented tumor expression profiles derived from SMOTE along with the source and test (unseen) data of melanoma patients treated with anti-PD1 therapy.
- **Figure S12.** t-SNE visualization of augmented microbiome profiles derived from DeepBioGen along with the source and test (unseen) data of diabetic patients and healthy controls.
- **Figure S13.** t-SNE visualization of augmented microbiome profiles derived from Random augmentation along with the source and test (unseen) data of diabetic patients and healthy controls.
- **Figure S14.** t-SNE visualization of augmented microbiome profiles derived from GMM along with the source and test (unseen) data of diabetic patients and healthy controls.
- **Figure S15.** t-SNE visualization of augmented microbiome profiles derived from SMOTE along with the source and test (unseen) data of diabetic patients and healthy controls.
- **Figure S16.** Feature-wise WSS by the number of clusters.
- **Figure S17.** Conditional Wasserstein GAN architecture in DeepBioGen.
- **Figure S18.** Sample-wise WSS by the number of clusters.

**Supplementary Tables:**

- **Table S1.** Summary of sequencing data sets.
- **Table S2.** Hyper-parameter grid for optimizing classifiers.
- **Table S3.** Modified inception scores of generated sequencing profiles varying the number of conditional Wasserstein GANs.
- **Table S4.** Anti-PD1 therapy responsiveness prediction accuracy of the best classifiers on unseen data.
- **Table S5.** Type 2 diabetes prediction accuracy of the best classifiers on unseen data.
- **Table S6.** 5-fold cross-validation on 126 tumor expression profiles (source data only).
- **Table S7.** 5-fold cross-validation on 344 gut microbiome profiles (source data only).
- **Table S8.** 5-fold cross-validation on 176 tumor expression profiles (source + test data).
- **Table S9.** 5-fold cross-validation on 440 gut microbiome profiles (source + test data).

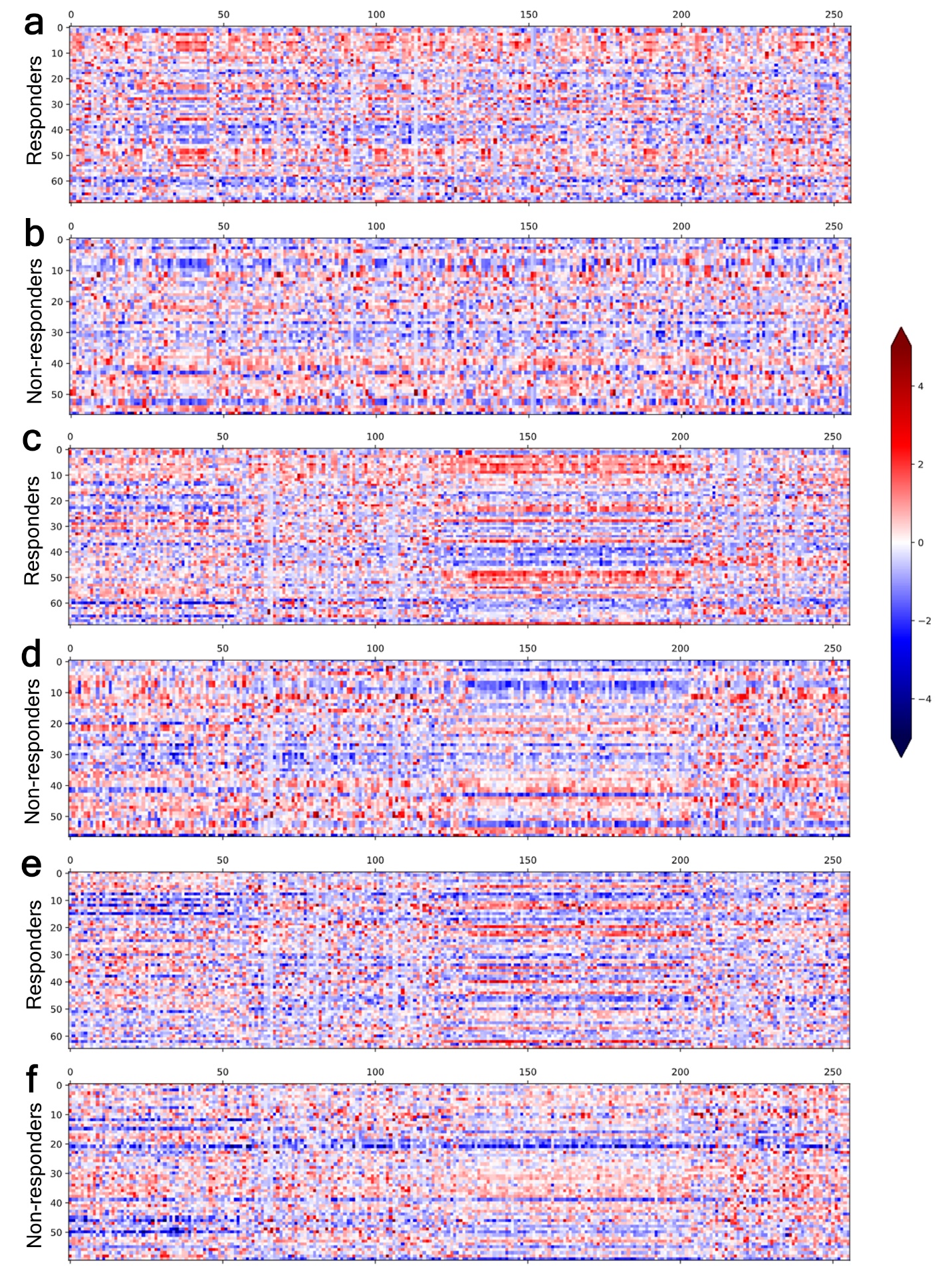
**Figure S1.** Visualization of tumor gene expression profiles of melanoma patients. **a-b**, The columns are unordered genes before pre-processing, and each row indicates the profile of responder (a) or non-responder (b). **c-d**, The columns are re-ordered genes with 4 clusters derived from feature-wise clustering. **e-f**, The augmented profiles generated by DeepBioGen.

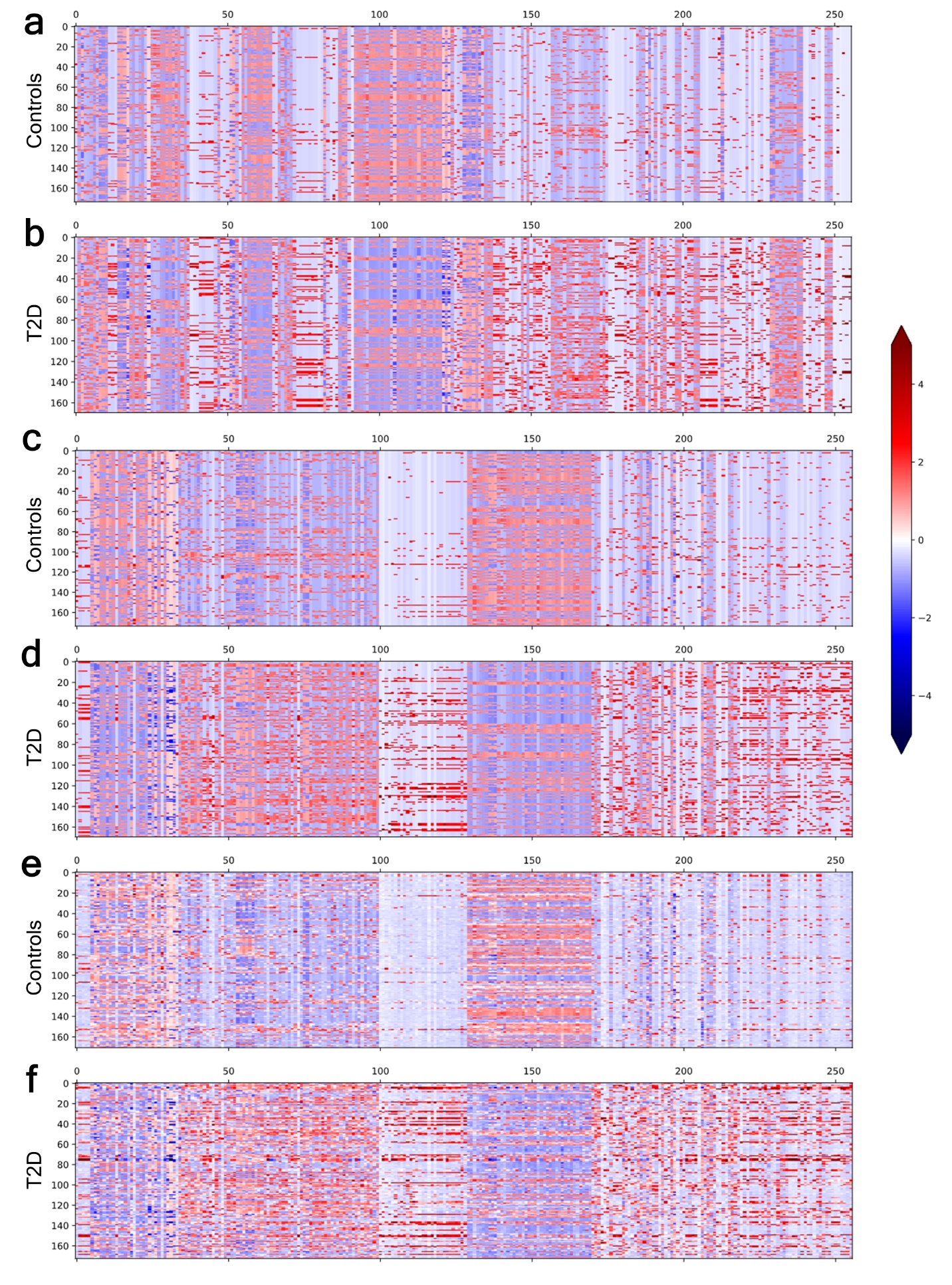
**Figure S2.** Visualization of microbiome marker profiles of diabetic patients and healthy controls. **a-b**, The columns are unordered genes before pre-processing, and each row indicates the profile of healthy controls (a) or type 2 diabetes (b). **c-d**, The columns are re-ordered genes with 4 clusters derived from feature-wise clustering. **e-f**, The augmented profiles generated by DeepBioGen.

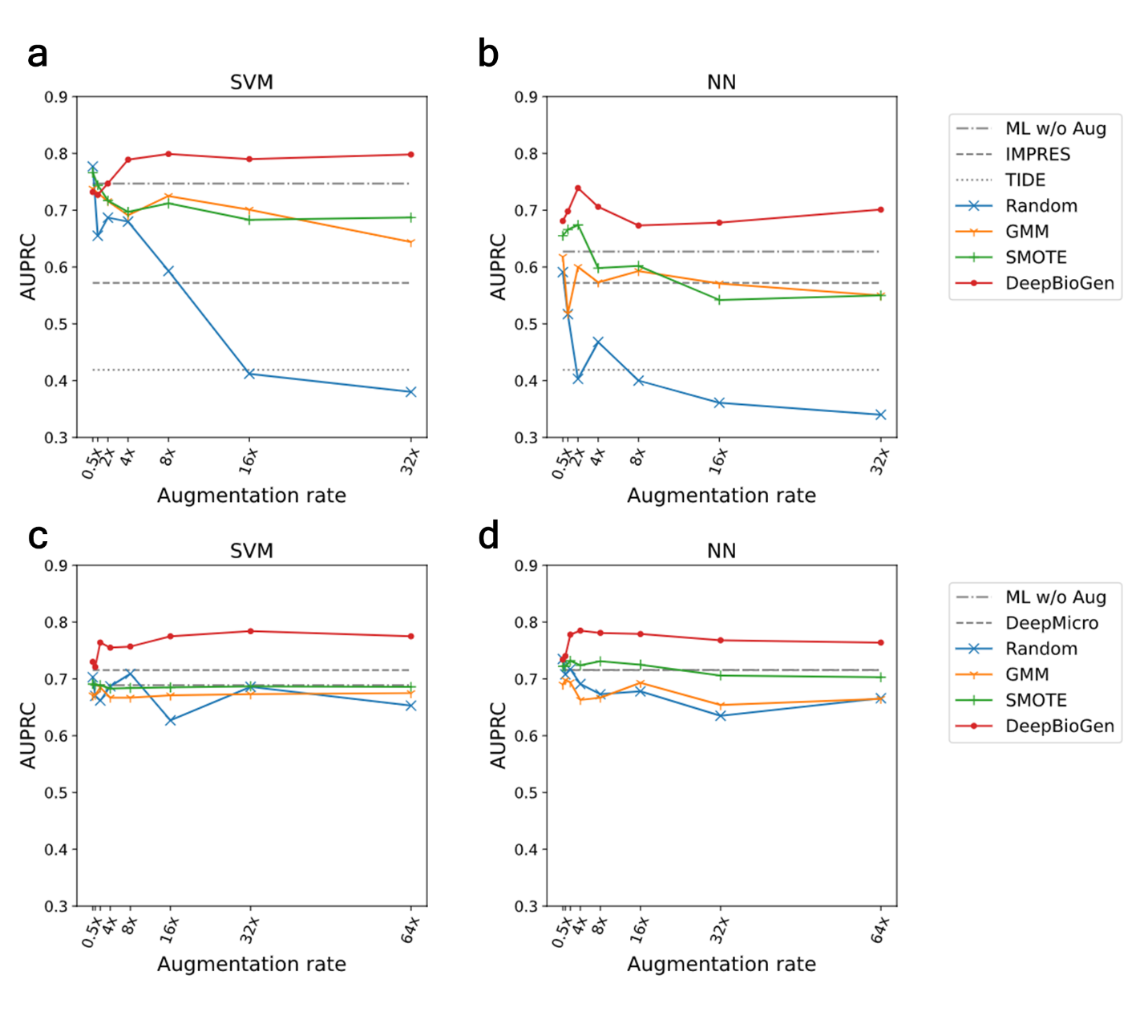

**Figure S3.** Prediction performance on unseen data (AUPRC). **a-b**, Results of anti-PD1 therapy response prediction on unseen data by the state-or-the-art and baseline classifiers (gray) and by classifiers generalized with DeepBioGen (red), SMOTE (green), GMM (yellow), and Random augmentation (blue); Classification algorithms: Support Vector Machine (SVM) and Neural network (NN) which is a multi-layer perceptron; Evaluation metric: Area under the precision-recall curve (AUPRC). **c-d**, Results of type 2 diabetes prediction on unseen data.

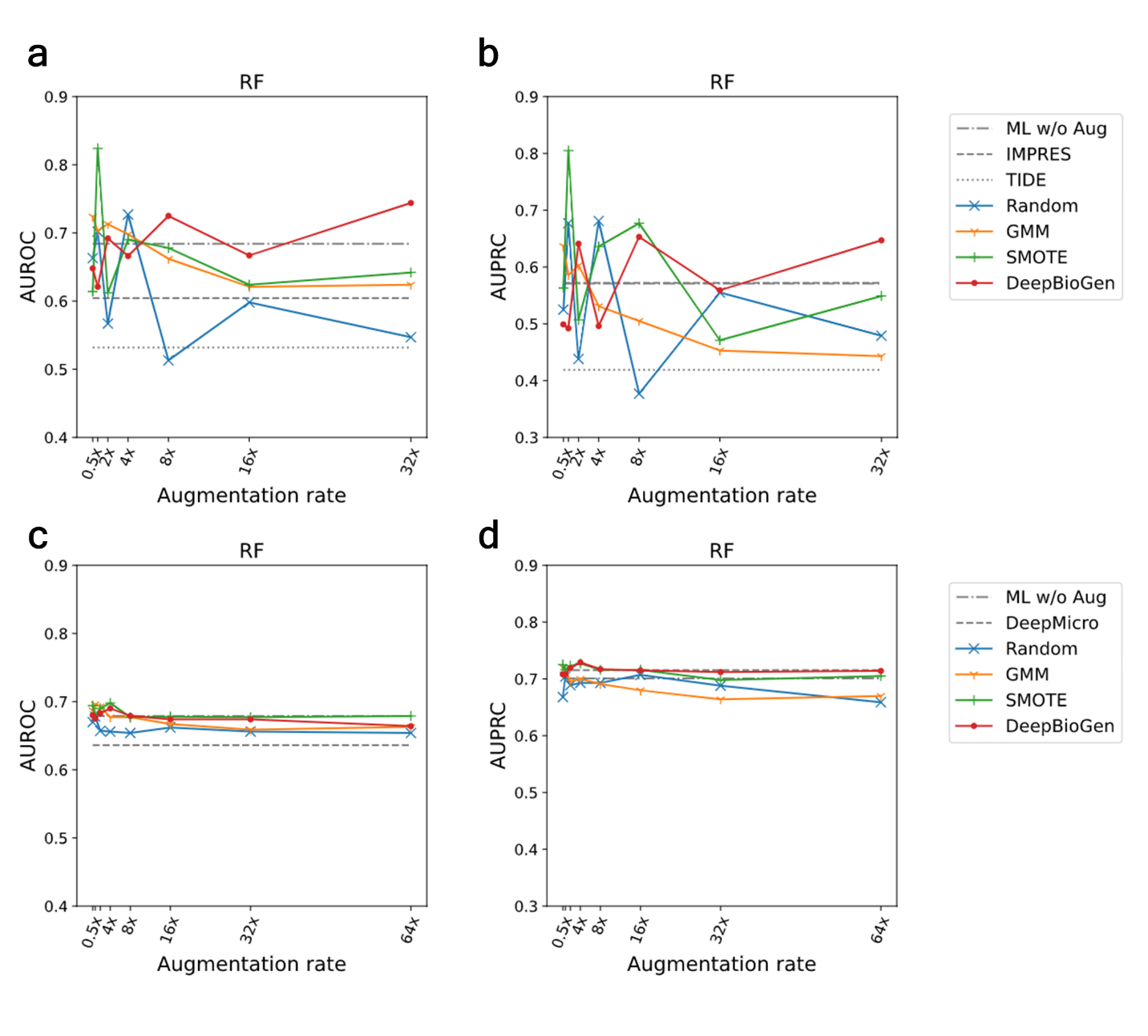

**Figure S4.** Prediction performance on unseen data with Random forest (RF) classifier. **a-b**, Results of anti-PD1 therapy response prediction on unseen data by the state-or-the-art and baseline classifiers (gray) and by the classifier generalized with DeepBioGen (red), SMOTE (green), GMM (yellow), and Random augmentation (blue); Evaluation metrics: Area under the receiver operating characteristics (AUROC) and Area under the precision-recall curve (AUPRC). **c-d**, Results of type 2 diabetes prediction on unseen data; Evaluation metric: AUROC and AUPRC.

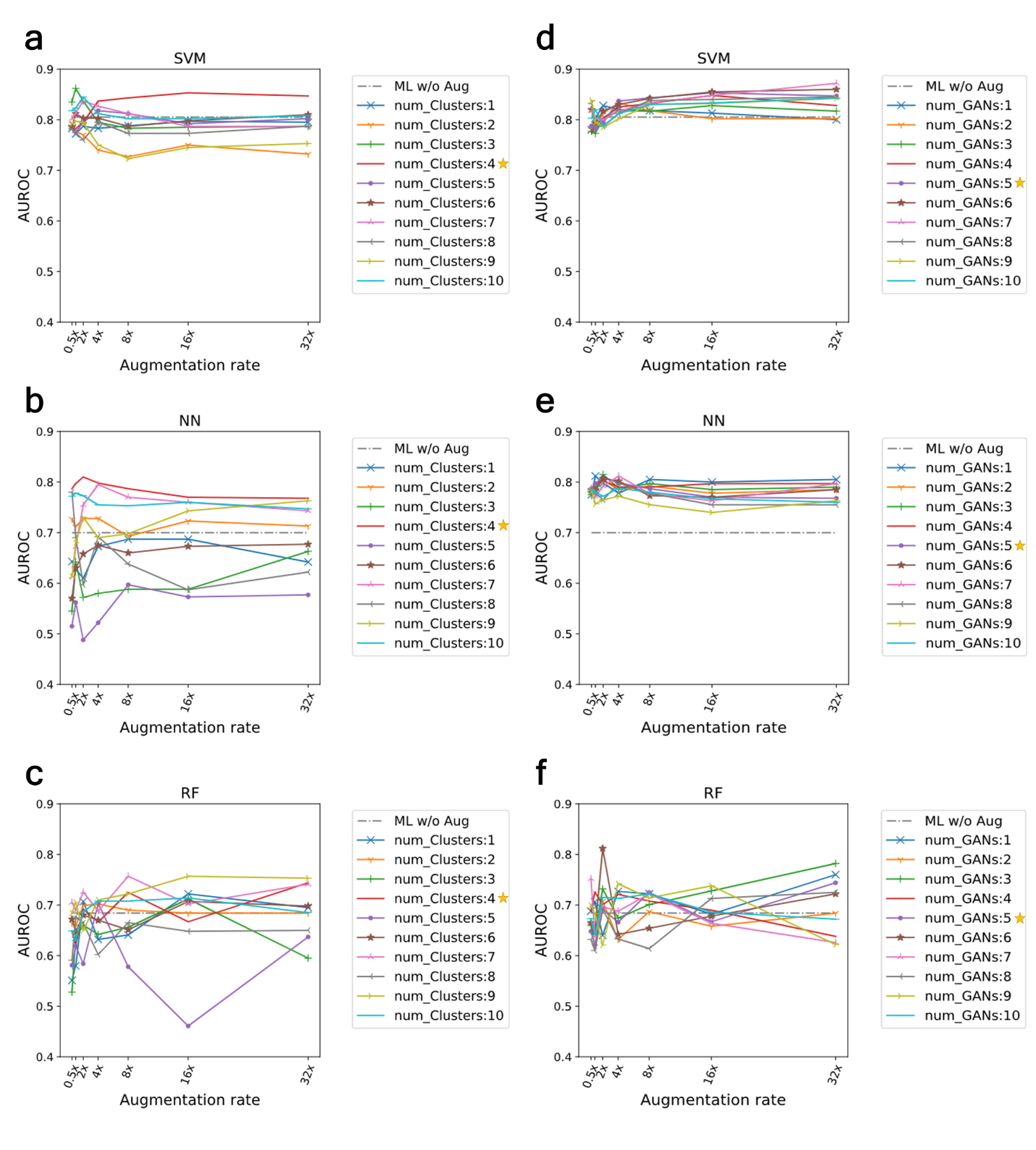

**Figure S5.** Anti-PD1 therapy response prediction performance (AUROC) on unseen data by varying the number of visual clusters and that of GAN models. **a-c**, Varying the number of visual clusters while fixing the number of GANs as five; Yellow star denotes the estimated number of visual clusters. **d-f**, Varying the number of GANs while fixing the number of visual clusters as four; Yellow start denotes the estimated number of GANs.

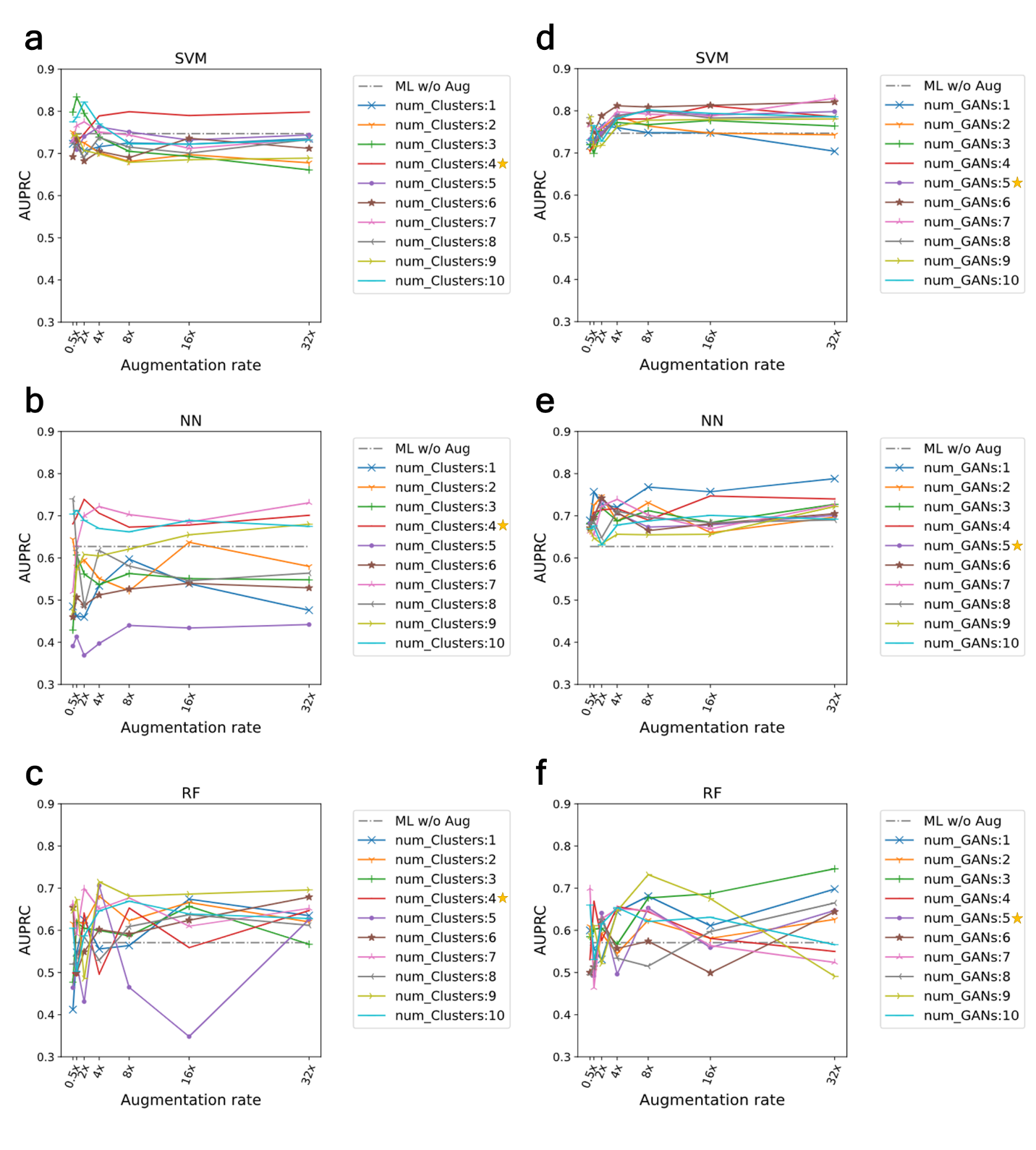

**Figure S6.** Anti-PD1 therapy response prediction performance (AUPRC) on unseen data by varying the number of visual clusters and that of GAN models. **a-c**, Varying the number of visual clusters while fixing the number of GANs as five; Yellow star denotes the estimated number of visual clusters. **d-f**, Varying the number of GANs while fixing the number of visual clusters as four; Yellow start denotes the estimated number of GANs.

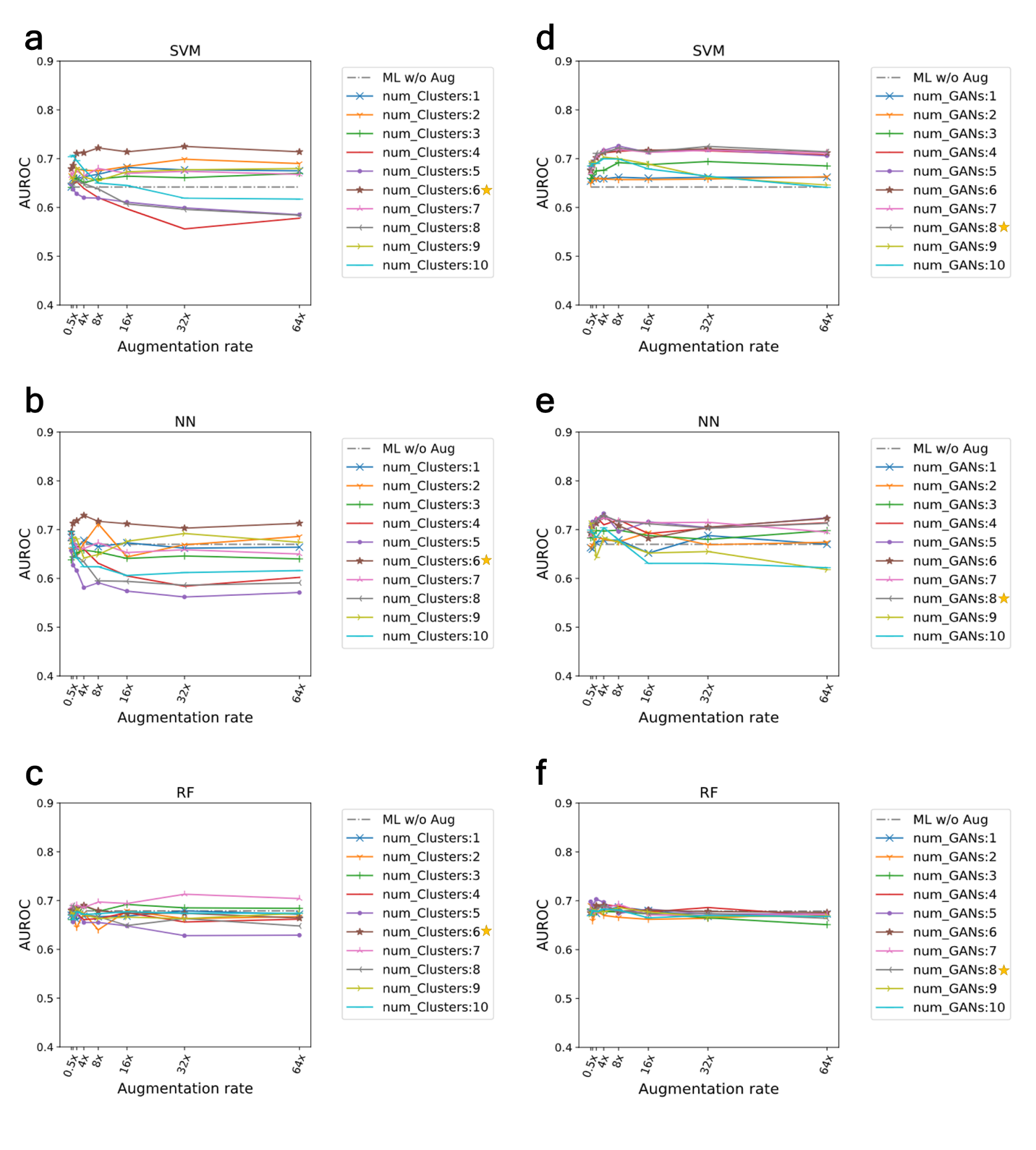

**Figure S7.** Type 2 diabetes prediction performance (AUROC) on unseen data by varying the number of visual clusters and that of GAN models. **a-c**, Varying the number of visual clusters while fixing the number of GANs as eight; Yellow star denotes the estimated number of visual clusters. **d-f**, Varying the number of GANs while fixing the number of visual clusters as six; Yellow start denotes the estimated number of GANs.

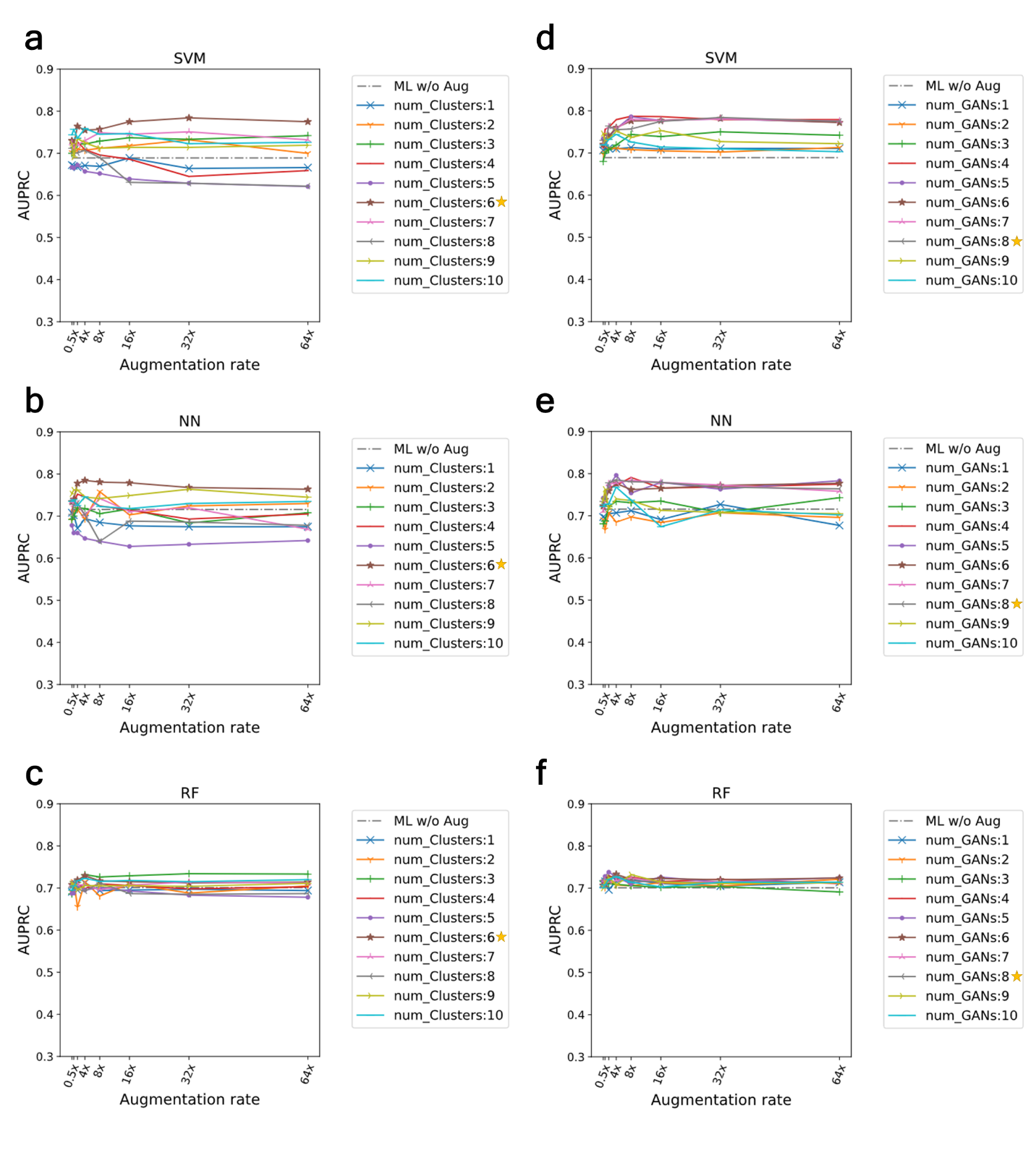

**Figure S8.** Type 2 diabetes prediction performance (AUPRC) on unseen data by varying the number of visual clusters and that of GAN models. **a-c**, Varying the number of visual clusters while fixing the number of GANs as eight; Yellow star denotes the estimated number of visual clusters. **d-f**, Varying the number of GANs while fixing the number of visual clusters as six; Yellow start denotes the estimated number of GANs.

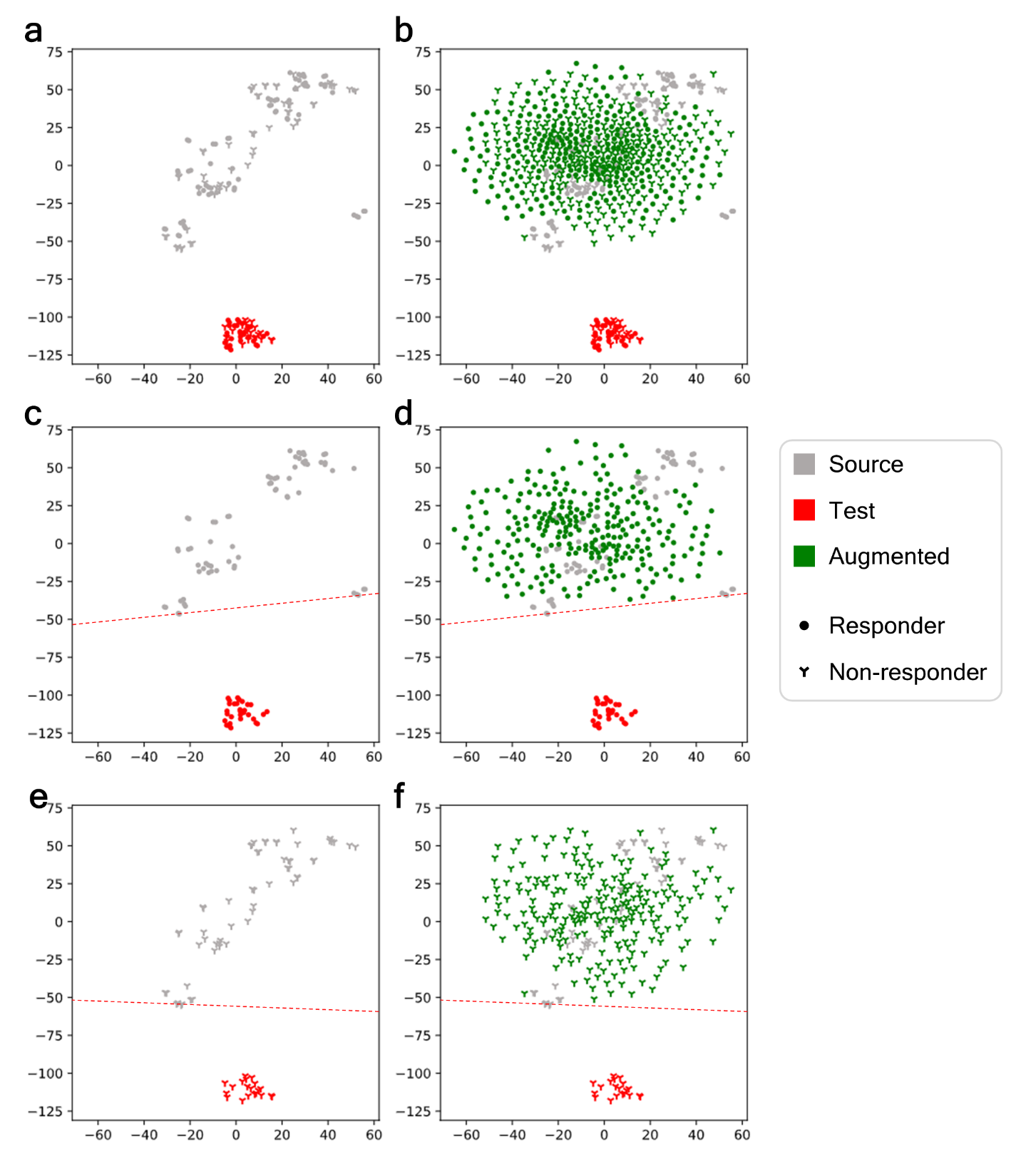

**Figure S9.** t-SNE visualization of augmented tumor expression profiles derived from Random augmentation along with the source and test (unseen) data of melanoma patients treated with anti-PD1 therapy. **a**, The source (gray) and test data (red). **b**, The source, test, and augmented data (green). **c**, Responders of the source and test data; An empirical boundary of responders of source data (red dotted line). **d**, Responders of the source, test, and augmented data. **e**, Non-responders of the source and test data; An empirical boundary of non-responders of source data (red dotted line). **f**, Non-responders of the source, test, and augmented data.

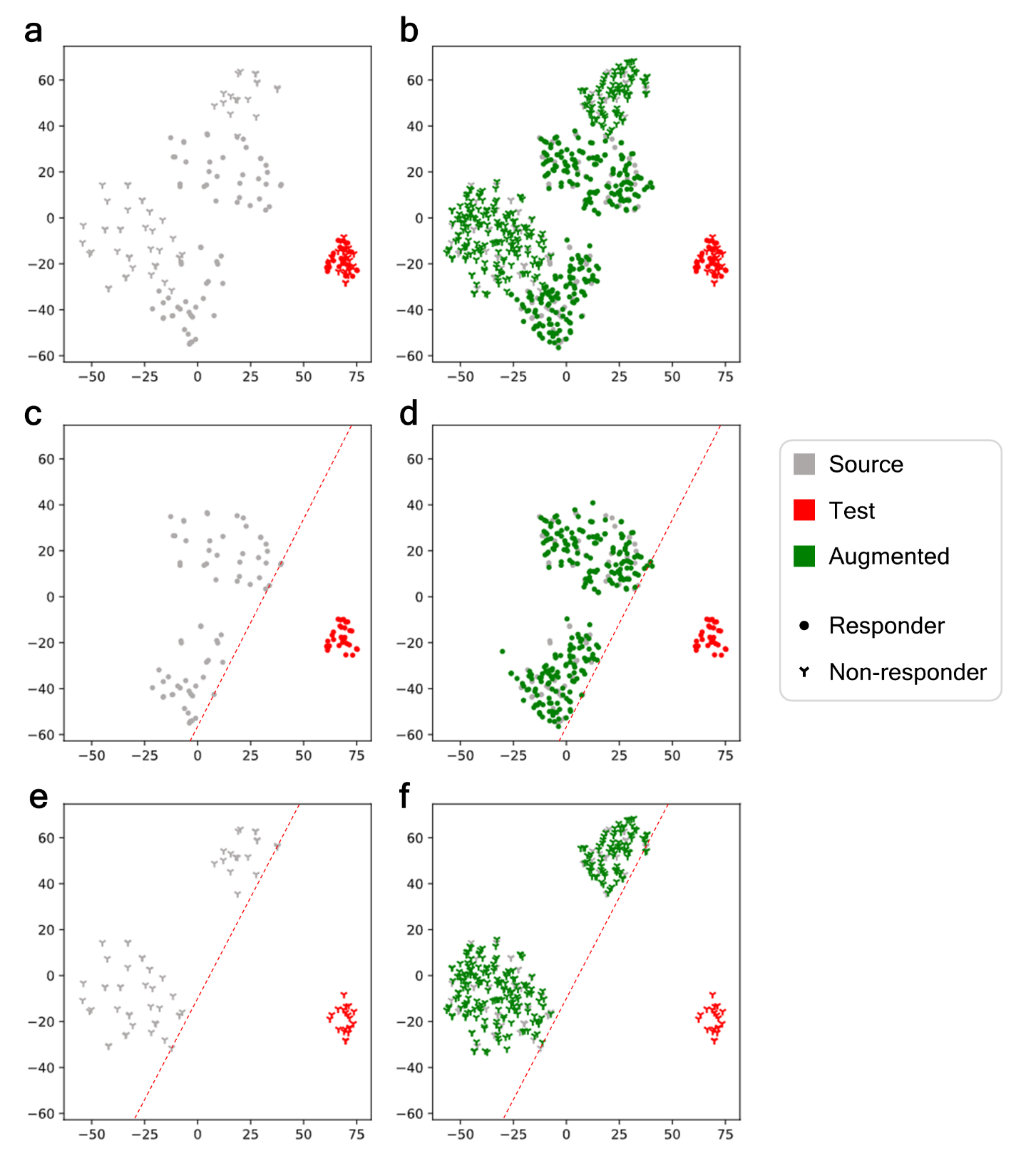

**Figure S10.** t-SNE visualization of augmented tumor expression profiles derived from GMM along with the source and test (unseen) data of melanoma patients treated with anti-PD1 therapy. **a**, The source (gray) and test data (red). **b**, The source, test, and augmented data (green). **c**, Responders of the source and test data; An empirical boundary of responders of source data (red dotted line). **d**, Responders of the source, test, and augmented data. **e**, Non-responders of the source and test data; An empirical boundary of non-responders of source data (red dotted line). **f**, Non-responders of the source, test, and augmented data.

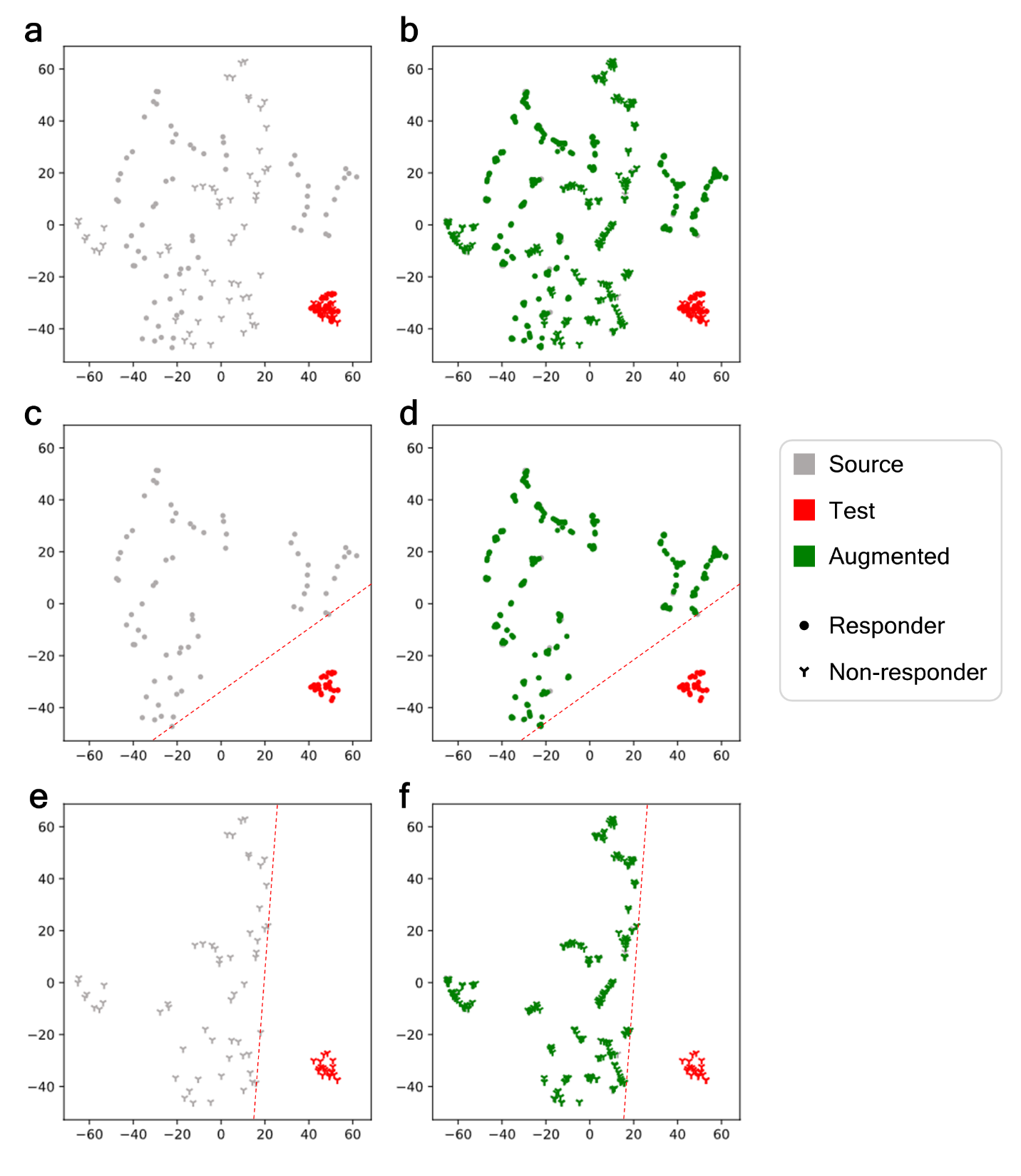

**Figure S11.** t-SNE visualization of augmented tumor expression profiles derived from SMOTE along with the source and test (unseen) data of melanoma patients treated with anti-PD1 therapy. **a**, The source (gray) and test data (red). **b**, The source, test, and augmented data (green). **c**, Responders of the source and test data; An empirical boundary of responders of source data (red dotted line). **d**, Responders of the source, test, and augmented data. **e**, Non-responders of the source and test data; An empirical boundary of non-responders of source data (red dotted line). **f**, Non-responders of the source, test, and augmented data.

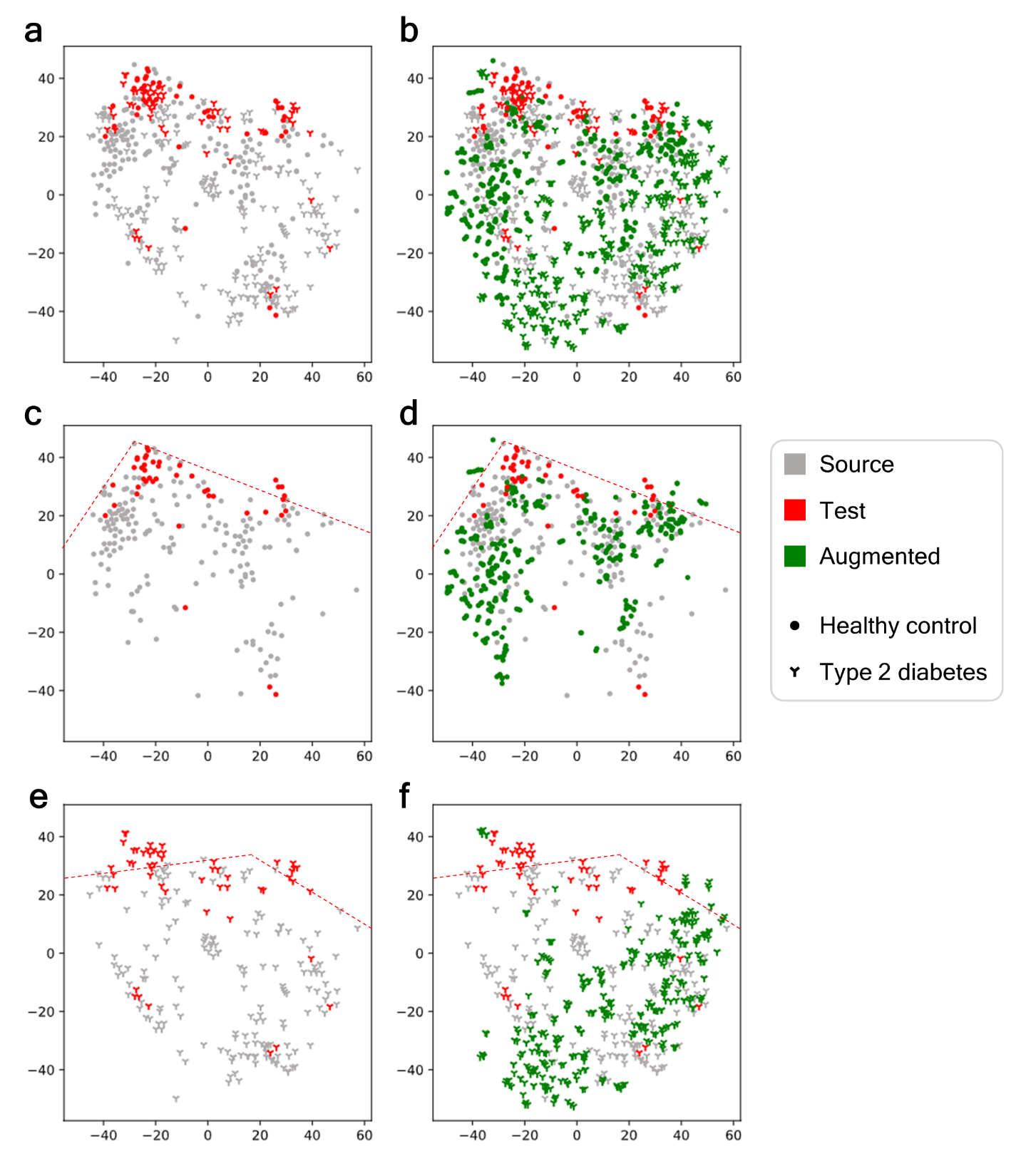

**Figure S12.** t-SNE visualization of augmented microbiome profiles derived from DeepBioGen along with the source and test (unseen) data of diabetic patients and healthy controls. **a**, The source (gray) and test data (red). **b**, The source, test, and augmented data (green). **c**, Healthy controls of the source and test data; An empirical boundary of healthy controls of source data (red dotted line). **d**, Healthy controls of the source, test, and augmented data. **e**, Type 2 diabetes patients of the source and test data; An empirical boundary of type 2 diabetes patients of source data (red dotted line). **f**, Type 2 diabetes patients of the source, test, and augmented data.

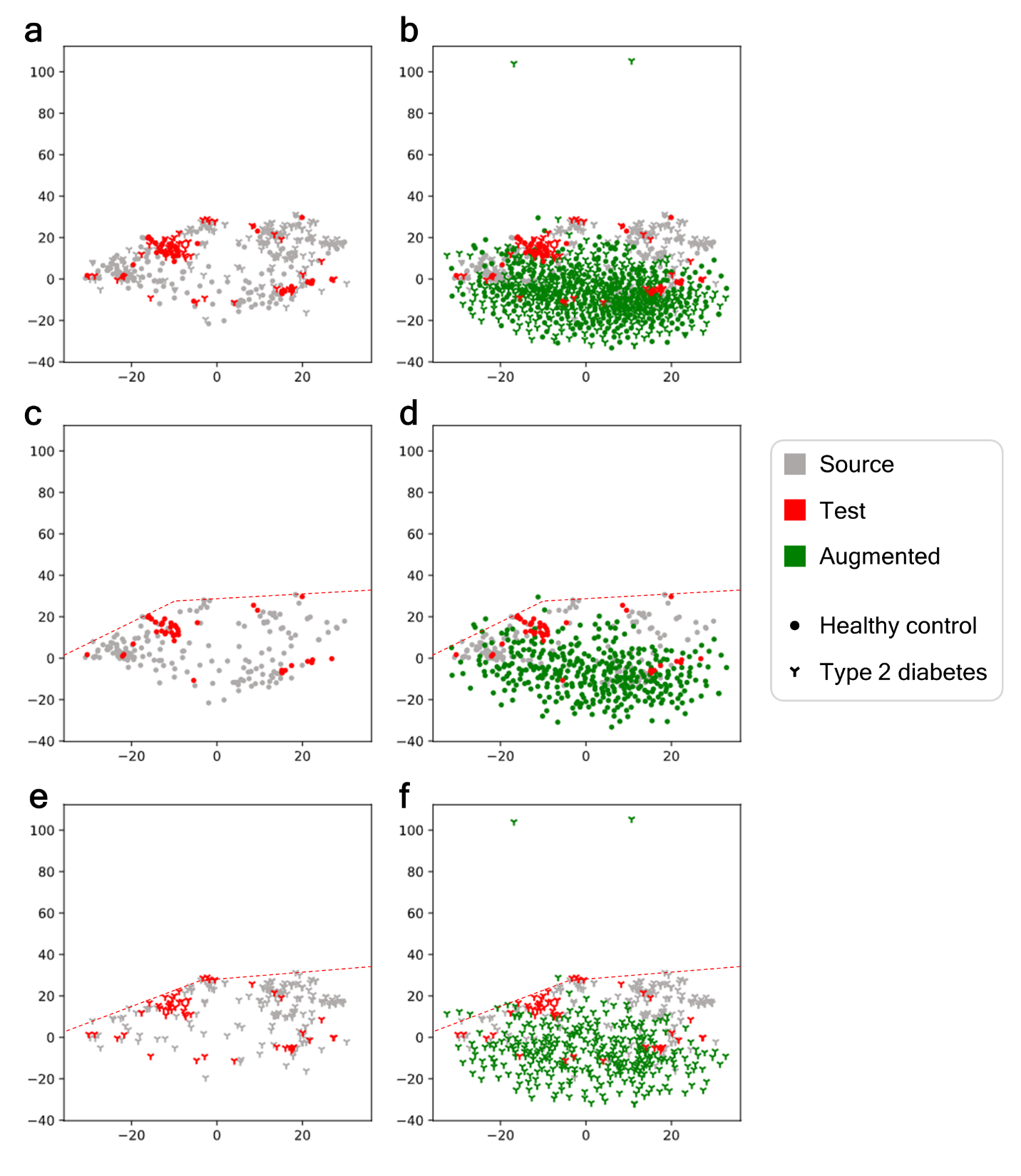

**Figure S13.** t-SNE visualization of augmented microbiome profiles derived from Random augmentation along with the source and test (unseen) data of diabetic patients and healthy controls. **a**, The source (gray) and test data (red). **b**, The source, test, and augmented data (green). **c**, Healthy controls of the source and test data; An empirical boundary of healthy controls of source data (red dotted line). **d**, Healthy controls of the source, test, and augmented data. **e**, Type 2 diabetes patients of the source and test data; An empirical boundary of type 2 diabetes patients of source data (red dotted line). **f**, Type 2 diabetes patients of the source, test, and augmented data.

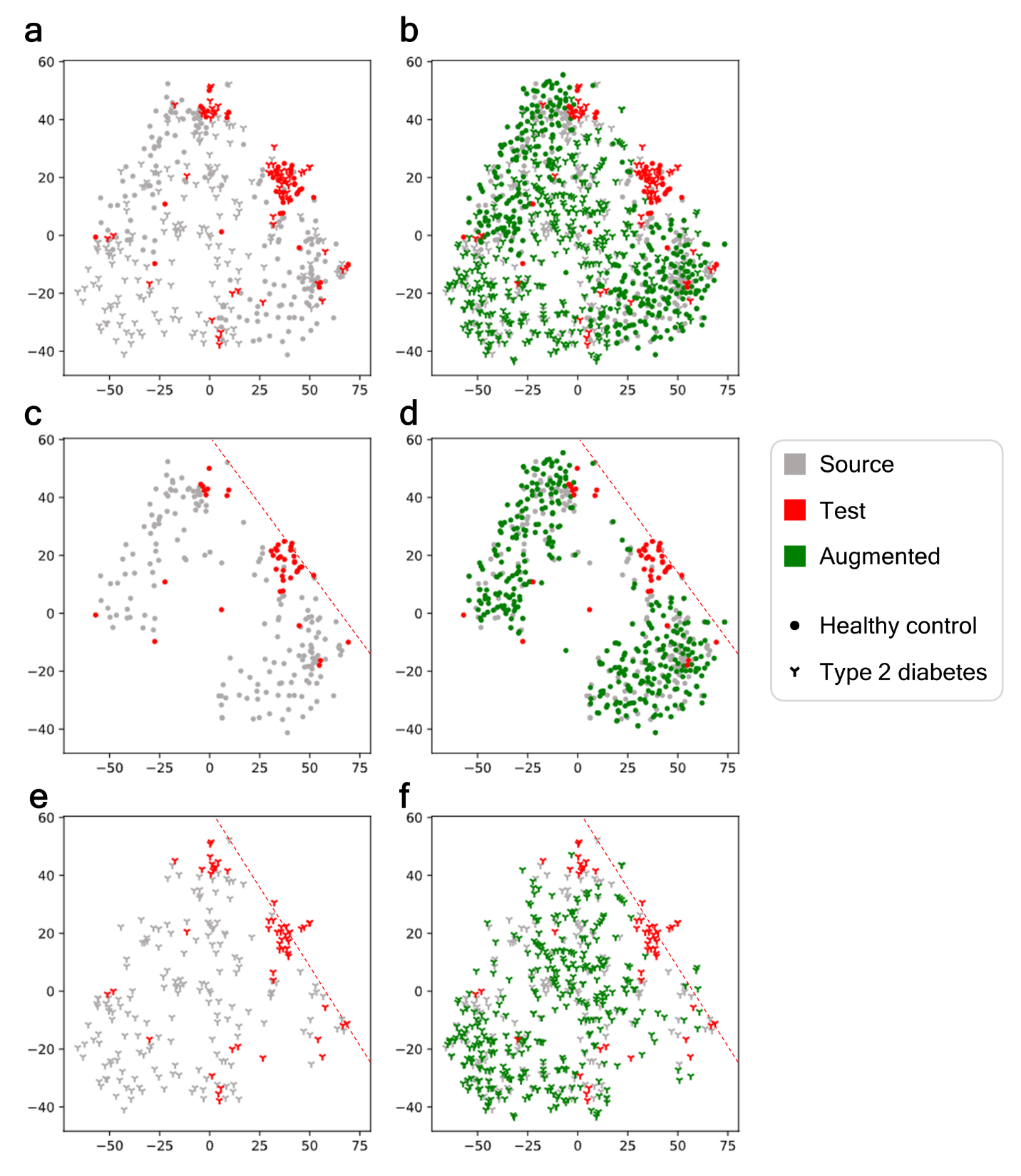

**Figure S14.** t-SNE visualization of augmented microbiome profiles derived from GMM along with the source and test (unseen) data of diabetic patients and healthy controls. **a**, The source (gray) and test data (red). **b**, The source, test, and augmented data (green). **c**, Healthy controls of the source and test data; An empirical boundary of healthy controls of source data (red dotted line). **d**, Healthy controls of the source, test, and augmented data. **e**, Type 2 diabetes patients of the source and test data; An empirical boundary of type 2 diabetes patients of source data (red dotted line). **f**, Type 2 diabetes patients of the source, test, and augmented data.

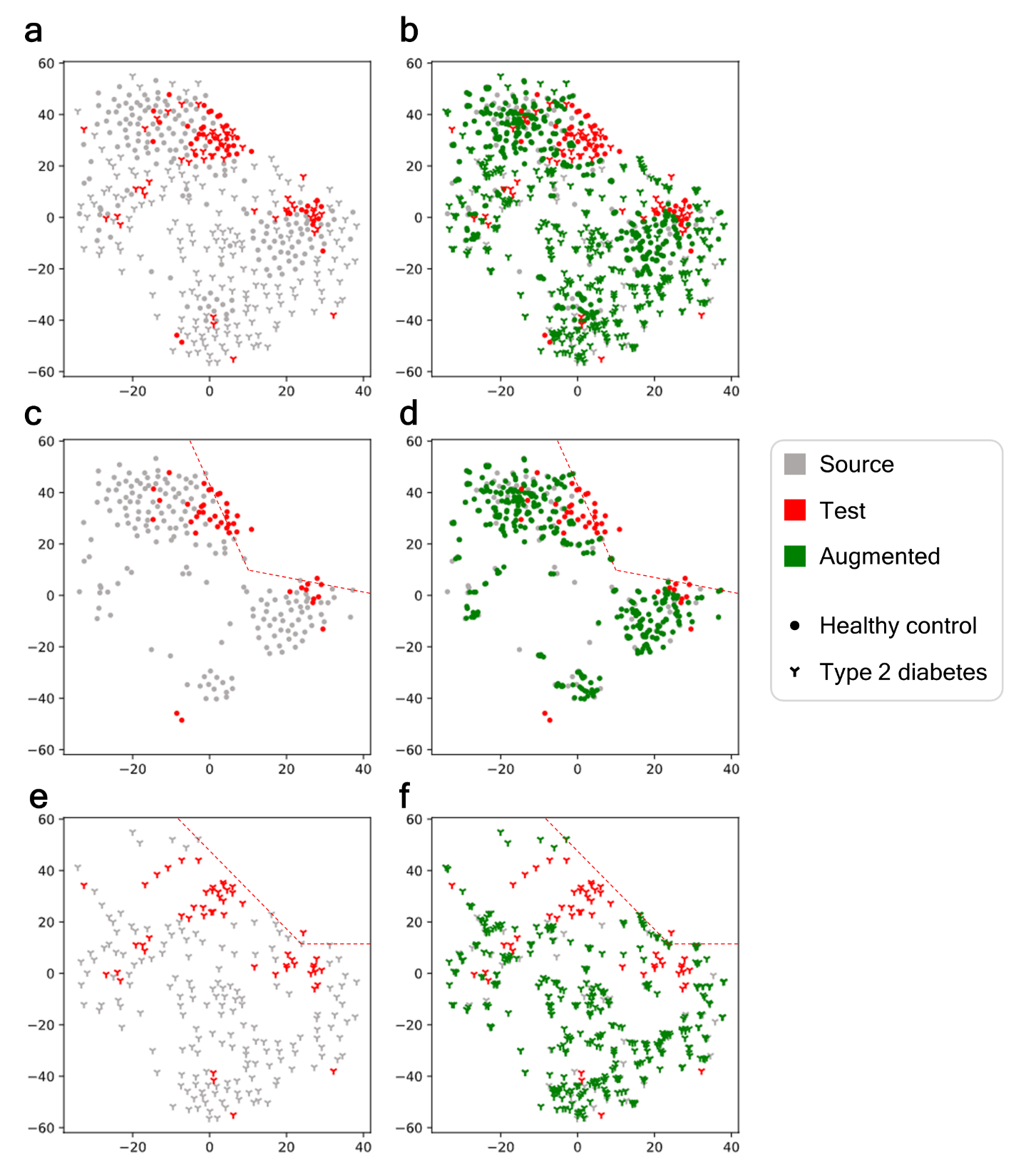

**Figure S15.** t-SNE visualization of augmented microbiome profiles derived from SMOTE along with the source and test (unseen) data of diabetic patients and healthy controls. **a**, The source (gray) and test data (red). **b**, The source, test, and augmented data (green). **c**, Healthy controls of the source and test data; An empirical boundary of healthy controls of source data (red dotted line). **d**, Healthy controls of the source, test, and augmented data. **e**, Type 2 diabetes patients of the source and test data; An empirical boundary of type 2 diabetes patients of source data (red dotted line). **f**, Type 2 diabetes patients of the source, test, and augmented data.

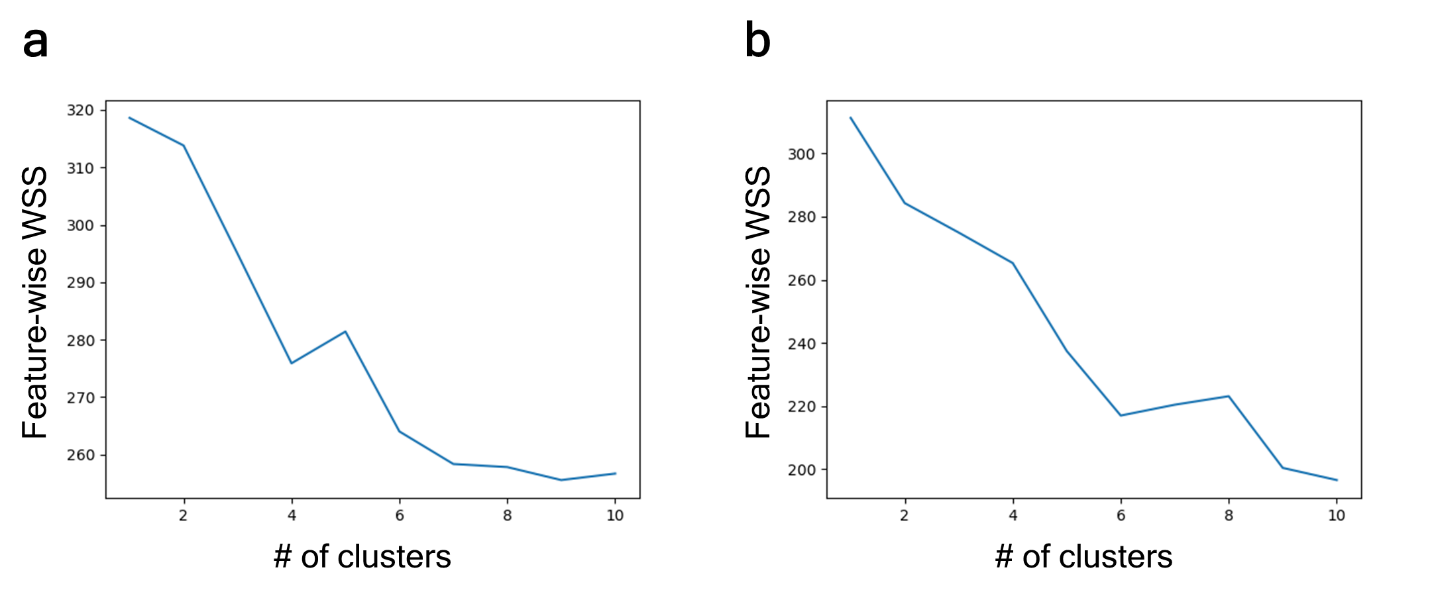

**Figure S16.** Feature-wise WSS by the number of clusters. **a**, RNA-seq tumor expression profiles (optimum: 4). **b**, WGS human gut microbiome marker profile (optimum: 6).

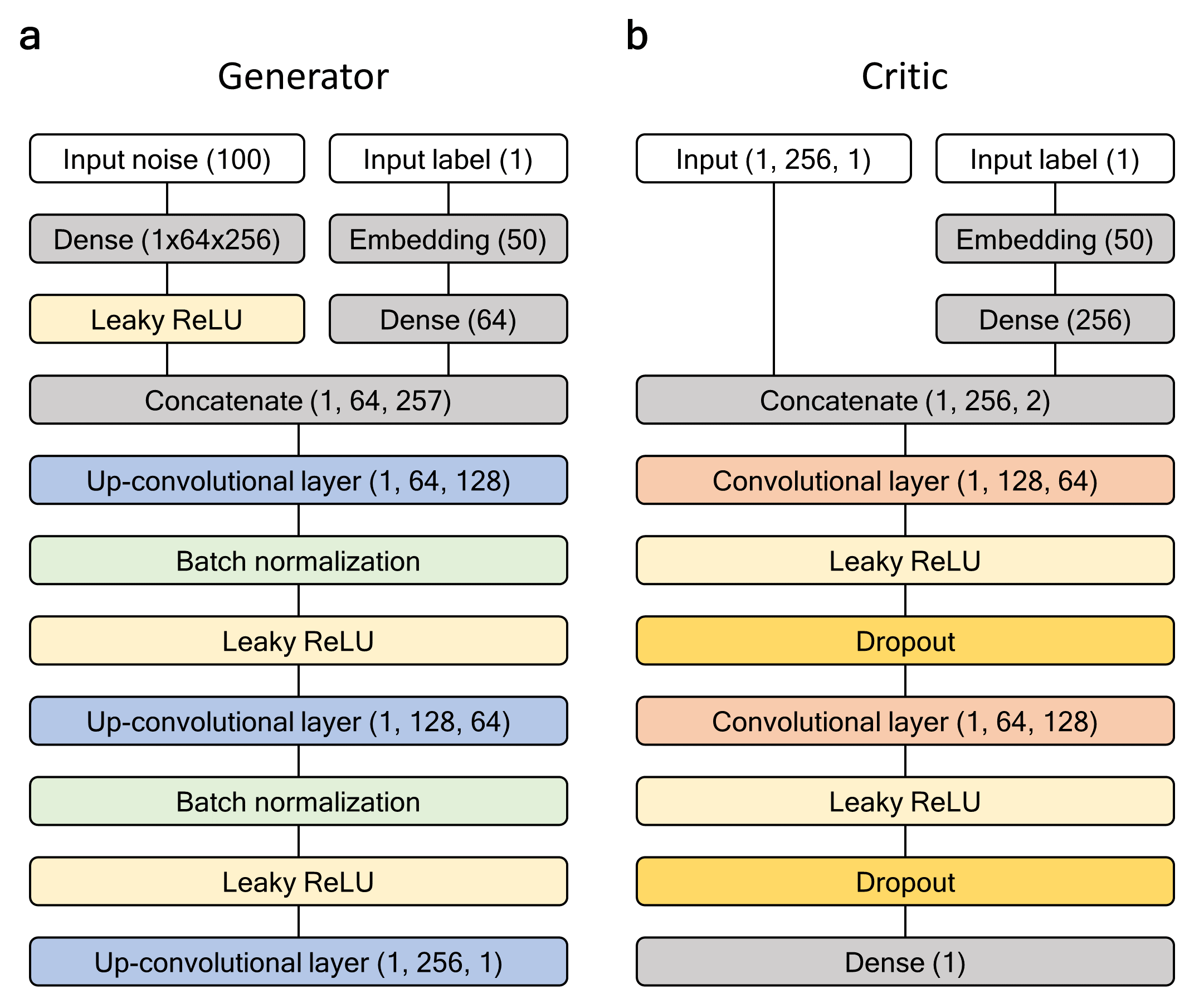

**Figure S17.** Conditional Wasserstein GAN architecture in DeepBioGen. **a**, A generator network generates realistic profiles from random noise. **b**, A critic network distinguishes realistic profiles from the real.

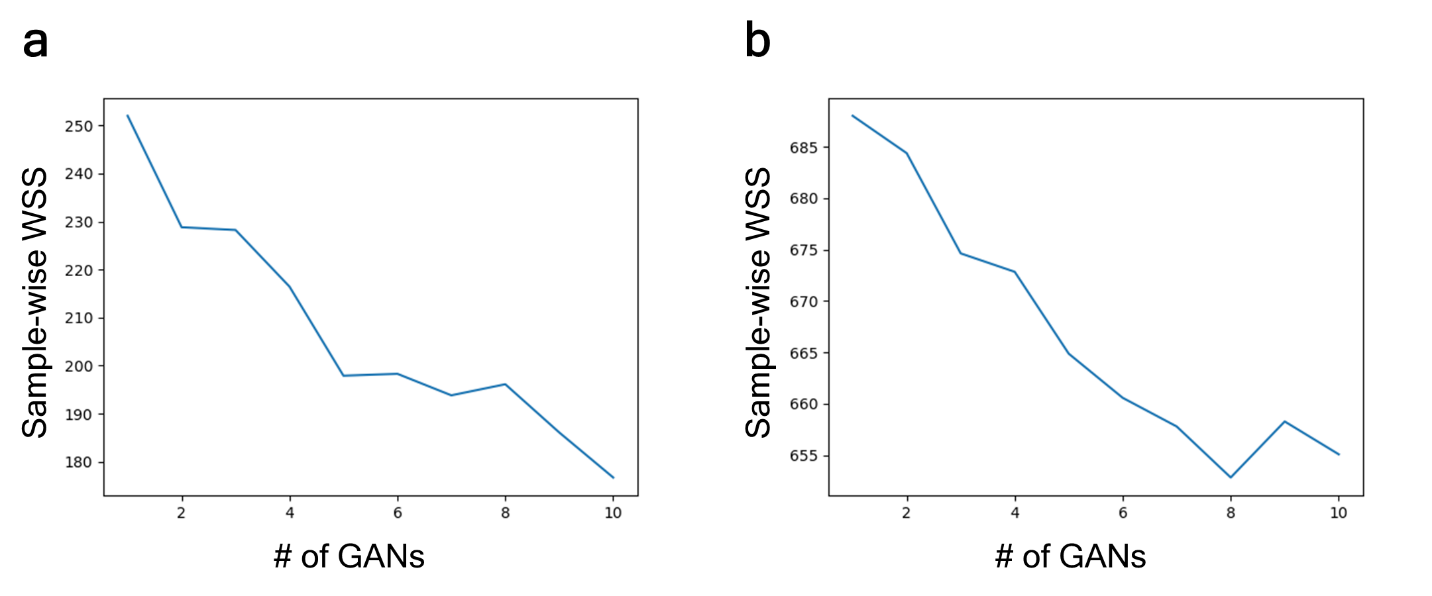
**Figure S18.** Sample-wise WSS by the number of GANs. **a**, RNA-seq tumor expression profiles (optimal: 5). **b**, WGS human gut microbiome marker profile (optimal: 8).

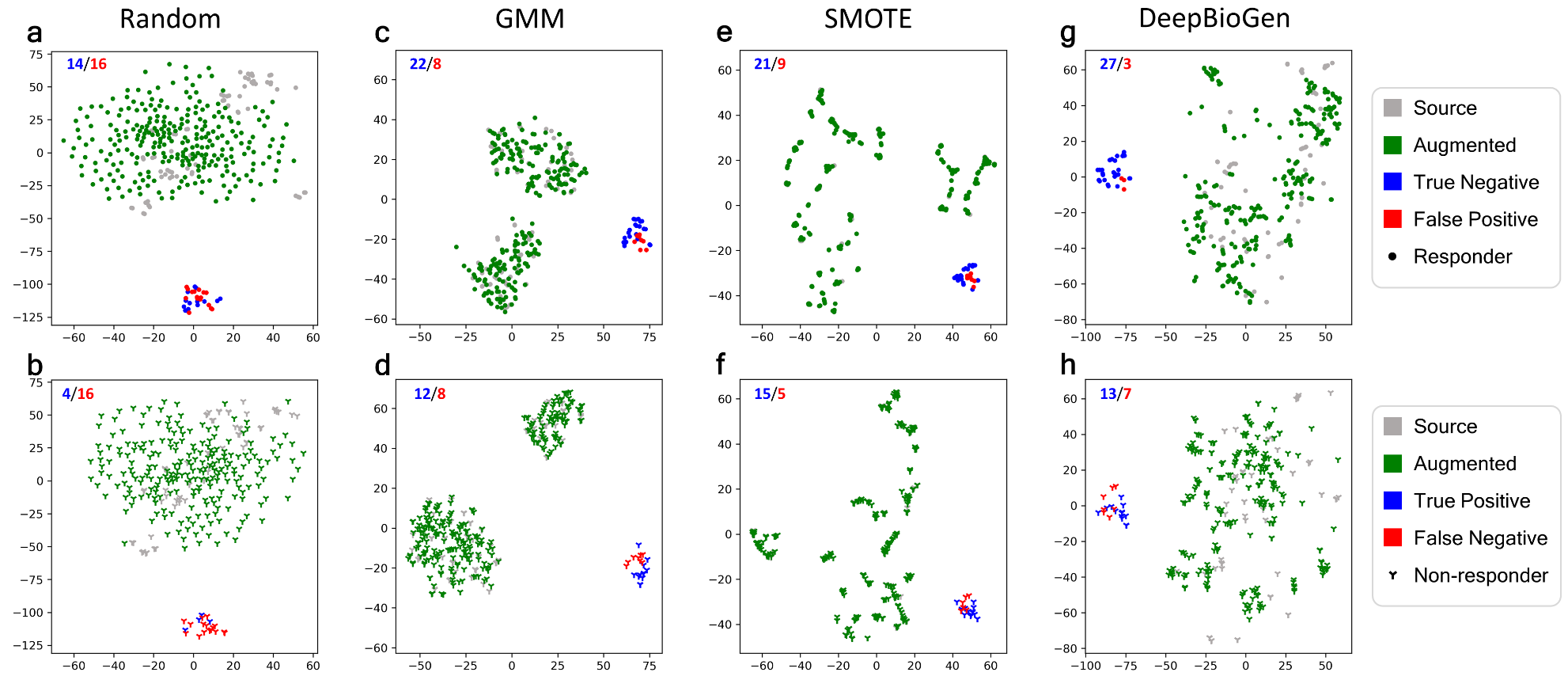

**Figure S19.** t-SNE visualization of anti-PD1 therapy responsiveness predictions on test data. **a-b**, predictions derived from SVM classifier based on random augmentation. **c-d**, predictions derived from SVM classifier based on GMM augmentation. **e-f**, predictions derived from SVM classifier based on SMOTE augmentation. **g-h**, predictions derived from SVM classifier based on DeepBioGen augmentation.

**Table S1.** Summary of sequencing data sets.

| Data type | | Year | # of Samples | # of class 0^*^ | # of class 1^**^ | Sequencing platform | # of original features | Reference |
| --- | --- | --- | --- | --- | --- | --- | --- | --- |
| RNA-seq tumor expression profile | Source | 2016 | 28 | 15 | 13 | Illumina Hiseq 2000 | 25,268 | Hugo et al.^1^ |
|  |  | 2017 | 98 | 54 | 44 | Illumina Hiseq 2000/2500 | 22,187 | Riaz et al.^2^ |
|  | Test | 2019 | 50 | 30 | 20 | Illumina Hiseq 2500 | 35,678 | Gide et al.^3^ |
| WGS human gut microbiome profile | Source | 2012 | 344 | 174 | 170 | Illumina Genome Analyzer II | 119,792 | Qin et al.^4^ |
|  | Test | 2014 | 96 | 43 | 53 | Illumina HiSeq 2000 | 83,456 | Karlsson et al.^5^ |

*Responders of anti-PD1 therapy for tumor expression profile or healthy controls for microbiome profile

**Non-responders of anti-PD1 therapy or type 2 diabetes for microbiome profile

**Table S2.** Hyper-parameter grid for optimizing classifiers.

| Classification algorithm | Hyper-parameter | Parameter grid |
| --- | --- | --- |
| SVM | Kernel | Linear and radial basis function (RBF) |
|  | Regularization penalty *C* | 2^-4^, 2^-3^, 2^-2^, 2^-1^, 2^0^, 2^1^, 2^2^, and 2^4^ |
|  | Gamma | ‘Scale’ (= 1/(n_features*X.var()) and  ‘Auto’ (=1/n_features) |
| RF | # of estimators | 2^7^, 2^8^, 2^9^, and 2^10^ |
|  | Maximum # of features for the best split | Square root and log2 of n_features |
|  | Split criterion | Gini impurity and information gain |
| NN | Hidden layers  (hidden units) | 3 layers (128, 64, 32),  4 layers (128, 64, 32, 16), and  5 layers (128, 64, 32, 16, 8) |
|  | Learning rate | Constant (0.001),  invscaling (0.001/ pow(t, power_t) where t is time step), and  adaptive (keep learning rate as long as training loss is decreasing, otherwise divide the current learning rate by 5) |
|  | Alpha (L2 penalty) | 0.0001, 0.001, 0.01, and 0.1 |

**Table S3.** Modified inception scores of generated sequencing profiles varying the number of conditional Wasserstein GANs.

| # of GANs | RNA-seq tumor expression profile | WGS human gut microbiome profile |
| --- | --- | --- |
| 1 | 1.0764 | 1.1923 |
| 2 | 1.0745 | 1.2046 |
| 3 | 1.0745 | 1.2079 |
| 4 | 1.0746 | 1.2048 |
| 5 | 1.0779 | 1.2060 |
| 6 | 1.0780 | 1.2061 |
| 7 | 1.0771 | 1.2071 |
| 8 | 1.0779 | 1.2042 |
| 9 | 1.0778 | 1.2031 |
| 10 | 1.0761 | 1.2035 |

Table S4. Anti-PD1 therapy responsiveness prediction accuracy of the best classifiers on unseen data.

| Augmentation Method | Classifier | F1^*^ | REC | PRE | ACC | TN | FP | FN | TP |
| --- | --- | --- | --- | --- | --- | --- | --- | --- | --- |
| Random | SVM | 0.2 | 0.2 | 0.2 | 0.36 | 14 | 16 | 16 | 4 |
| Random | RF | 0.27 | 0.25 | 0.294 | 0.46 | 18 | 12 | 15 | 5 |
| Random | NN | 0.519 | 0.7 | 0.412 | 0.48 | 10 | 20 | 6 | 14 |
| GMM | SVM | 0.6 | 0.6 | 0.6 | 0.68 | 22 | 8 | 8 | 12 |
| GMM | RF | 0.605 | 0.65 | 0.565 | 0.66 | 20 | 10 | 7 | 13 |
| GMM | NN | 0.696 | **0.8** | 0.615 | 0.72 | 20 | 10 | 4 | 16 |
| Smote | SVM | 0.651 | 0.7 | 0.609 | 0.7 | 21 | 9 | 6 | 14 |
| Smote | RF | 0.536 | 0.75 | 0.417 | 0.48 | 9 | 21 | 5 | 15 |
| Smote | NN | 0.486 | 0.45 | 0.529 | 0.62 | 22 | 8 | 11 | 9 |
| DeepBioGen | SVM | **0.722^**^** | 0.65 | **0.812** | **0.8** | 27 | 3 | 7 | 13 |
| DeepBioGen | RF | 0.619 | 0.65 | 0.591 | 0.68 | 21 | 9 | 7 | 13 |
| DeepBioGen | NN | 0.7 | 0.7 | 0.7 | 0.76 | 24 | 6 | 6 | 14 |

*F1: F-measure (Harmonic mean of precision and recall); REC: Recall; PRE: Precision; ACC: Accuracy; TN: True Negative; FP: False Positive; FN: False Negative; TP: True Positive

**The bolded value indicates the best performance across the column

Table S5. Type 2 diabetes prediction accuracy of the best classifiers on unseen data.

| Augmentation Method | Classifier | F1^*^ | REC | PRE | ACC | TN | FP | FN | TP |
| --- | --- | --- | --- | --- | --- | --- | --- | --- | --- |
| Random | SVM | 0.511 | 0.434 | 0.622 | 0.542 | 29 | 14 | 30 | 23 |
| Random | RF | 0.475 | 0.453 | 0.5 | 0.448 | 19 | 24 | 29 | 24 |
| Random | NN | 0.521 | 0.472 | 0.581 | 0.521 | 25 | 18 | 28 | 25 |
| GMM | SVM | 0.707 | **0.774** | 0.651 | 0.646 | 21 | 22 | 12 | 41 |
| GMM | RF | 0.696 | 0.755 | 0.645 | 0.635 | 21 | 22 | 13 | 40 |
| GMM | NN | 0.706 | 0.679 | 0.735 | **0.688** | 30 | 13 | 17 | 36 |
| Smote | SVM | 0.697 | 0.717 | 0.679 | 0.656 | 25 | 18 | 15 | 38 |
| Smote | RF | 0.68 | 0.642 | 0.723 | 0.667 | 30 | 13 | 19 | 34 |
| Smote | NN | 0.7 | 0.66 | **0.745** | **0.688** | 31 | 12 | 18 | 35 |
| DeepBioGen | SVM | **0.712^**^** | 0.698 | 0.725 | **0.688** | 29 | 14 | 16 | 37 |
| DeepBioGen | RF | 0.624 | 0.547 | 0.725 | 0.635 | 32 | 11 | 24 | 29 |
| DeepBioGen | NN | 0.698 | 0.698 | 0.698 | 0.667 | 27 | 16 | 16 | 37 |

*F1: F-measure (Harmonic mean of precision and recall); REC: Recall; PRE: Precision; ACC: Accuracy; TN: True Negative; FP: False Positive; FN: False Negative; TP: True Positive

**The bolded value indicates the best performance across the column

Table S6. 5-fold cross-validation on 126 tumor expression profiles (source data only).

| Random Split | Clf | AUROC | STD | AUPRC | STD | ACC | STD | REC | STD | PRE | STD | F1 | STD |
| --- | --- | --- | --- | --- | --- | --- | --- | --- | --- | --- | --- | --- | --- |
| 0 | SVM | 0.811 | 0.075 | 0.806 | 0.086 | 0.738 | 0.043 | 0.702 | 0.066 | 0.716 | 0.111 | 0.704 | 0.064 |
| 1 | SVM | 0.791 | 0.034 | 0.725 | 0.126 | 0.699 | 0.052 | 0.685 | 0.164 | 0.664 | 0.103 | 0.657 | 0.104 |
| 2 | SVM | 0.766 | 0.087 | 0.735 | 0.111 | 0.707 | 0.095 | 0.621 | 0.172 | 0.726 | 0.116 | 0.646 | 0.112 |
| 3 | SVM | 0.712 | 0.100 | 0.658 | 0.181 | 0.620 | 0.078 | 0.636 | 0.122 | 0.597 | 0.129 | 0.598 | 0.075 |
| 4 | SVM | 0.783 | 0.114 | 0.783 | 0.121 | 0.722 | 0.083 | 0.647 | 0.141 | 0.726 | 0.069 | 0.674 | 0.093 |
| 5 | SVM | 0.707 | 0.041 | 0.642 | 0.165 | 0.603 | 0.020 | 0.577 | 0.114 | 0.594 | 0.146 | 0.557 | 0.037 |
| 6 | SVM | 0.692 | 0.086 | 0.660 | 0.123 | 0.570 | 0.081 | 0.466 | 0.084 | 0.580 | 0.225 | 0.490 | 0.084 |
| 7 | SVM | 0.749 | 0.106 | 0.680 | 0.185 | 0.626 | 0.095 | 0.579 | 0.083 | 0.628 | 0.214 | 0.583 | 0.100 |
| 8 | SVM | 0.702 | 0.087 | 0.652 | 0.117 | 0.610 | 0.114 | 0.510 | 0.232 | 0.574 | 0.131 | 0.523 | 0.162 |
| 9 | SVM | 0.673 | 0.068 | 0.622 | 0.201 | 0.603 | 0.100 | 0.447 | 0.128 | 0.557 | 0.201 | 0.491 | 0.149 |
| 0 | RF | 0.747 | 0.098 | 0.751 | 0.087 | 0.660 | 0.146 | 0.554 | 0.250 | 0.649 | 0.160 | 0.577 | 0.192 |
| 1 | RF | 0.712 | 0.092 | 0.677 | 0.093 | 0.636 | 0.091 | 0.561 | 0.107 | 0.646 | 0.162 | 0.580 | 0.067 |
| 2 | RF | 0.721 | 0.090 | 0.693 | 0.075 | 0.652 | 0.103 | 0.553 | 0.239 | 0.644 | 0.131 | 0.558 | 0.173 |
| 3 | RF | 0.693 | 0.047 | 0.638 | 0.100 | 0.643 | 0.026 | 0.606 | 0.177 | 0.641 | 0.147 | 0.591 | 0.072 |
| 4 | RF | 0.709 | 0.156 | 0.722 | 0.125 | 0.627 | 0.134 | 0.530 | 0.148 | 0.630 | 0.129 | 0.563 | 0.113 |
| 5 | RF | 0.670 | 0.055 | 0.615 | 0.157 | 0.571 | 0.051 | 0.471 | 0.151 | 0.556 | 0.136 | 0.479 | 0.106 |
| 6 | RF | 0.661 | 0.077 | 0.660 | 0.119 | 0.555 | 0.071 | 0.435 | 0.081 | 0.580 | 0.236 | 0.464 | 0.055 |
| 7 | RF | 0.733 | 0.099 | 0.698 | 0.152 | 0.658 | 0.085 | 0.537 | 0.114 | 0.729 | 0.241 | 0.582 | 0.091 |
| 8 | RF | 0.656 | 0.065 | 0.653 | 0.085 | 0.579 | 0.071 | 0.400 | 0.149 | 0.563 | 0.127 | 0.450 | 0.109 |
| 9 | RF | 0.686 | 0.097 | 0.678 | 0.188 | 0.642 | 0.069 | 0.447 | 0.128 | 0.650 | 0.273 | 0.511 | 0.161 |
| 0 | NN | 0.788 | 0.068 | 0.767 | 0.090 | 0.738 | 0.054 | 0.719 | 0.118 | 0.705 | 0.065 | 0.708 | 0.075 |
| 1 | NN | 0.718 | 0.083 | 0.621 | 0.168 | 0.674 | 0.049 | 0.640 | 0.124 | 0.637 | 0.125 | 0.630 | 0.102 |
| 2 | NN | 0.747 | 0.107 | 0.662 | 0.085 | 0.692 | 0.090 | 0.692 | 0.153 | 0.660 | 0.085 | 0.663 | 0.086 |
| 3 | NN | 0.702 | 0.057 | 0.625 | 0.157 | 0.643 | 0.081 | 0.583 | 0.085 | 0.638 | 0.132 | 0.597 | 0.063 |
| 4 | NN | 0.745 | 0.138 | 0.714 | 0.150 | 0.699 | 0.107 | 0.652 | 0.160 | 0.688 | 0.078 | 0.660 | 0.100 |
| 5 | NN | 0.689 | 0.094 | 0.627 | 0.198 | 0.587 | 0.053 | 0.608 | 0.071 | 0.562 | 0.142 | 0.565 | 0.060 |
| 6 | NN | 0.683 | 0.081 | 0.655 | 0.112 | 0.627 | 0.061 | 0.524 | 0.080 | 0.650 | 0.199 | 0.554 | 0.054 |
| 7 | NN | 0.714 | 0.103 | 0.633 | 0.168 | 0.666 | 0.085 | 0.631 | 0.041 | 0.650 | 0.172 | 0.631 | 0.091 |
| 8 | NN | 0.698 | 0.114 | 0.649 | 0.132 | 0.610 | 0.084 | 0.561 | 0.193 | 0.579 | 0.109 | 0.553 | 0.112 |
| 9 | NN | 0.653 | 0.050 | 0.572 | 0.180 | 0.659 | 0.060 | 0.558 | 0.131 | 0.602 | 0.127 | 0.577 | 0.126 |

Table S7. 5-fold cross-validation on 344 gut microbiome profiles (source data only).

| Random Split | Clf | AUROC | STD | AUPRC | STD | ACC | STD | REC | STD | PRE | STD | F1 | STD |
| --- | --- | --- | --- | --- | --- | --- | --- | --- | --- | --- | --- | --- | --- |
| 0 | SVM | 0.727 | 0.022 | 0.699 | 0.043 | 0.683 | 0.026 | 0.707 | 0.035 | 0.672 | 0.060 | 0.686 | 0.029 |
| 1 | SVM | 0.715 | 0.097 | 0.704 | 0.072 | 0.657 | 0.101 | 0.699 | 0.103 | 0.646 | 0.076 | 0.670 | 0.083 |
| 2 | SVM | 0.731 | 0.049 | 0.715 | 0.074 | 0.686 | 0.059 | 0.702 | 0.053 | 0.683 | 0.095 | 0.688 | 0.054 |
| 3 | SVM | 0.729 | 0.033 | 0.702 | 0.052 | 0.669 | 0.027 | 0.709 | 0.076 | 0.655 | 0.048 | 0.676 | 0.023 |
| 4 | SVM | 0.726 | 0.046 | 0.713 | 0.057 | 0.678 | 0.056 | 0.697 | 0.072 | 0.668 | 0.067 | 0.680 | 0.057 |
| 5 | SVM | 0.728 | 0.044 | 0.699 | 0.074 | 0.683 | 0.039 | 0.688 | 0.041 | 0.676 | 0.080 | 0.679 | 0.051 |
| 6 | SVM | 0.731 | 0.097 | 0.719 | 0.106 | 0.677 | 0.098 | 0.728 | 0.102 | 0.663 | 0.099 | 0.690 | 0.087 |
| 7 | SVM | 0.726 | 0.041 | 0.703 | 0.045 | 0.672 | 0.042 | 0.731 | 0.097 | 0.654 | 0.036 | 0.685 | 0.040 |
| 8 | SVM | 0.750 | 0.032 | 0.737 | 0.066 | 0.694 | 0.036 | 0.693 | 0.078 | 0.697 | 0.078 | 0.689 | 0.045 |
| 9 | SVM | 0.745 | 0.034 | 0.741 | 0.054 | 0.648 | 0.051 | 0.663 | 0.082 | 0.647 | 0.105 | 0.645 | 0.062 |
| 0 | RF | 0.726 | 0.014 | 0.701 | 0.041 | 0.645 | 0.036 | 0.602 | 0.027 | 0.659 | 0.071 | 0.626 | 0.022 |
| 1 | RF | 0.712 | 0.113 | 0.698 | 0.082 | 0.660 | 0.104 | 0.623 | 0.103 | 0.675 | 0.092 | 0.646 | 0.093 |
| 2 | RF | 0.710 | 0.026 | 0.694 | 0.050 | 0.657 | 0.030 | 0.621 | 0.075 | 0.664 | 0.059 | 0.638 | 0.049 |
| 3 | RF | 0.728 | 0.049 | 0.707 | 0.059 | 0.666 | 0.076 | 0.616 | 0.094 | 0.681 | 0.115 | 0.642 | 0.090 |
| 4 | RF | 0.728 | 0.028 | 0.692 | 0.062 | 0.657 | 0.034 | 0.616 | 0.054 | 0.668 | 0.032 | 0.639 | 0.026 |
| 5 | RF | 0.721 | 0.046 | 0.691 | 0.071 | 0.640 | 0.039 | 0.624 | 0.073 | 0.647 | 0.091 | 0.628 | 0.044 |
| 6 | RF | 0.720 | 0.102 | 0.716 | 0.113 | 0.660 | 0.097 | 0.673 | 0.117 | 0.650 | 0.090 | 0.660 | 0.097 |
| 7 | RF | 0.728 | 0.029 | 0.705 | 0.047 | 0.651 | 0.036 | 0.666 | 0.101 | 0.647 | 0.030 | 0.650 | 0.043 |
| 8 | RF | 0.709 | 0.055 | 0.703 | 0.106 | 0.654 | 0.018 | 0.636 | 0.082 | 0.664 | 0.069 | 0.642 | 0.032 |
| 9 | RF | 0.721 | 0.050 | 0.708 | 0.066 | 0.642 | 0.045 | 0.611 | 0.064 | 0.658 | 0.095 | 0.624 | 0.031 |
| 0 | NN | 0.726 | 0.025 | 0.698 | 0.035 | 0.651 | 0.040 | 0.640 | 0.061 | 0.651 | 0.080 | 0.642 | 0.054 |
| 1 | NN | 0.709 | 0.086 | 0.704 | 0.065 | 0.678 | 0.081 | 0.674 | 0.068 | 0.682 | 0.070 | 0.676 | 0.061 |
| 2 | NN | 0.711 | 0.058 | 0.687 | 0.099 | 0.663 | 0.056 | 0.684 | 0.077 | 0.658 | 0.095 | 0.665 | 0.061 |
| 3 | NN | 0.714 | 0.044 | 0.694 | 0.065 | 0.654 | 0.063 | 0.641 | 0.122 | 0.654 | 0.065 | 0.641 | 0.077 |
| 4 | NN | 0.727 | 0.054 | 0.712 | 0.078 | 0.654 | 0.051 | 0.628 | 0.070 | 0.659 | 0.049 | 0.641 | 0.050 |
| 5 | NN | 0.727 | 0.043 | 0.692 | 0.081 | 0.660 | 0.031 | 0.644 | 0.043 | 0.664 | 0.087 | 0.649 | 0.042 |
| 6 | NN | 0.728 | 0.096 | 0.711 | 0.108 | 0.662 | 0.101 | 0.686 | 0.097 | 0.656 | 0.099 | 0.668 | 0.090 |
| 7 | NN | 0.725 | 0.031 | 0.697 | 0.044 | 0.646 | 0.051 | 0.657 | 0.142 | 0.642 | 0.029 | 0.639 | 0.076 |
| 8 | NN | 0.754 | 0.041 | 0.745 | 0.052 | 0.671 | 0.054 | 0.631 | 0.104 | 0.680 | 0.086 | 0.650 | 0.077 |
| 9 | NN | 0.738 | 0.052 | 0.746 | 0.041 | 0.663 | 0.055 | 0.641 | 0.102 | 0.674 | 0.064 | 0.648 | 0.036 |

Table S8. 5-fold cross-validation on 176 tumor expression profiles (source + test data).

| Random Split | Clf | AUROC | STD | AUPRC | STD | ACC | STD | REC | STD | PRE | STD | F1 | STD |
| --- | --- | --- | --- | --- | --- | --- | --- | --- | --- | --- | --- | --- | --- |
| 0 | SVM | 0.738 | 0.106 | 0.705 | 0.089 | 0.682 | 0.087 | 0.663 | 0.099 | 0.639 | 0.083 | 0.646 | 0.073 |
| 1 | SVM | 0.747 | 0.095 | 0.690 | 0.138 | 0.659 | 0.090 | 0.598 | 0.123 | 0.623 | 0.173 | 0.599 | 0.124 |
| 2 | SVM | 0.691 | 0.087 | 0.651 | 0.081 | 0.626 | 0.095 | 0.620 | 0.211 | 0.584 | 0.125 | 0.579 | 0.113 |
| 3 | SVM | 0.734 | 0.108 | 0.685 | 0.100 | 0.665 | 0.092 | 0.631 | 0.154 | 0.637 | 0.105 | 0.617 | 0.097 |
| 4 | SVM | 0.709 | 0.056 | 0.654 | 0.008 | 0.630 | 0.065 | 0.590 | 0.090 | 0.586 | 0.108 | 0.579 | 0.078 |
| 5 | SVM | 0.745 | 0.098 | 0.641 | 0.170 | 0.710 | 0.098 | 0.604 | 0.180 | 0.685 | 0.146 | 0.633 | 0.145 |
| 6 | SVM | 0.744 | 0.086 | 0.670 | 0.147 | 0.705 | 0.069 | 0.607 | 0.087 | 0.690 | 0.154 | 0.641 | 0.103 |
| 7 | SVM | 0.734 | 0.083 | 0.645 | 0.187 | 0.653 | 0.082 | 0.582 | 0.123 | 0.642 | 0.209 | 0.586 | 0.100 |
| 8 | SVM | 0.723 | 0.102 | 0.655 | 0.155 | 0.636 | 0.075 | 0.536 | 0.161 | 0.599 | 0.178 | 0.549 | 0.142 |
| 9 | SVM | 0.720 | 0.052 | 0.685 | 0.096 | 0.647 | 0.031 | 0.549 | 0.137 | 0.619 | 0.078 | 0.567 | 0.073 |
| 0 | RF | 0.709 | 0.111 | 0.704 | 0.092 | 0.631 | 0.147 | 0.567 | 0.223 | 0.576 | 0.105 | 0.559 | 0.168 |
| 1 | RF | 0.717 | 0.124 | 0.671 | 0.188 | 0.630 | 0.102 | 0.463 | 0.073 | 0.643 | 0.240 | 0.524 | 0.125 |
| 2 | RF | 0.640 | 0.102 | 0.604 | 0.116 | 0.574 | 0.070 | 0.446 | 0.192 | 0.524 | 0.133 | 0.459 | 0.126 |
| 3 | RF | 0.709 | 0.113 | 0.688 | 0.119 | 0.636 | 0.057 | 0.516 | 0.103 | 0.610 | 0.126 | 0.549 | 0.076 |
| 4 | RF | 0.675 | 0.065 | 0.617 | 0.076 | 0.557 | 0.086 | 0.477 | 0.122 | 0.511 | 0.132 | 0.480 | 0.089 |
| 5 | RF | 0.706 | 0.098 | 0.656 | 0.154 | 0.648 | 0.092 | 0.489 | 0.128 | 0.621 | 0.169 | 0.543 | 0.141 |
| 6 | RF | 0.728 | 0.088 | 0.671 | 0.163 | 0.648 | 0.065 | 0.497 | 0.127 | 0.641 | 0.138 | 0.546 | 0.096 |
| 7 | RF | 0.702 | 0.087 | 0.638 | 0.174 | 0.636 | 0.085 | 0.537 | 0.128 | 0.629 | 0.196 | 0.555 | 0.080 |
| 8 | RF | 0.703 | 0.067 | 0.643 | 0.109 | 0.665 | 0.085 | 0.559 | 0.091 | 0.646 | 0.139 | 0.591 | 0.088 |
| 9 | RF | 0.702 | 0.023 | 0.650 | 0.054 | 0.637 | 0.059 | 0.471 | 0.196 | 0.624 | 0.123 | 0.510 | 0.137 |
| 0 | NN | 0.732 | 0.130 | 0.695 | 0.125 | 0.694 | 0.101 | 0.707 | 0.163 | 0.643 | 0.120 | 0.663 | 0.114 |
| 1 | NN | 0.717 | 0.108 | 0.638 | 0.197 | 0.670 | 0.093 | 0.623 | 0.120 | 0.615 | 0.179 | 0.614 | 0.140 |
| 2 | NN | 0.638 | 0.094 | 0.571 | 0.093 | 0.626 | 0.080 | 0.577 | 0.159 | 0.592 | 0.120 | 0.566 | 0.090 |
| 3 | NN | 0.717 | 0.080 | 0.630 | 0.145 | 0.664 | 0.080 | 0.600 | 0.120 | 0.628 | 0.074 | 0.606 | 0.081 |
| 4 | NN | 0.693 | 0.071 | 0.593 | 0.082 | 0.618 | 0.099 | 0.611 | 0.157 | 0.568 | 0.110 | 0.575 | 0.118 |
| 5 | NN | 0.720 | 0.119 | 0.646 | 0.181 | 0.711 | 0.122 | 0.562 | 0.197 | 0.702 | 0.200 | 0.615 | 0.185 |
| 6 | NN | 0.731 | 0.091 | 0.648 | 0.153 | 0.688 | 0.087 | 0.637 | 0.096 | 0.661 | 0.157 | 0.641 | 0.103 |
| 7 | NN | 0.738 | 0.092 | 0.637 | 0.164 | 0.687 | 0.033 | 0.640 | 0.091 | 0.659 | 0.127 | 0.633 | 0.032 |
| 8 | NN | 0.726 | 0.099 | 0.632 | 0.183 | 0.653 | 0.073 | 0.563 | 0.133 | 0.608 | 0.153 | 0.577 | 0.122 |
| 9 | NN | 0.705 | 0.043 | 0.636 | 0.056 | 0.631 | 0.042 | 0.501 | 0.121 | 0.616 | 0.090 | 0.534 | 0.065 |

Table S9. 5-fold cross-validation on 440 gut microbiome profiles (source + test data).

| Random Split | Clf | AUROC | STD | AUPRC | STD | ACC | STD | REC | STD | PRE | STD | F1 | STD |
| --- | --- | --- | --- | --- | --- | --- | --- | --- | --- | --- | --- | --- | --- |
| 0 | SVM | 0.723 | 0.057 | 0.733 | 0.061 | 0.636 | 0.069 | 0.691 | 0.082 | 0.628 | 0.058 | 0.657 | 0.066 |
| 1 | SVM | 0.740 | 0.029 | 0.755 | 0.053 | 0.679 | 0.035 | 0.751 | 0.057 | 0.662 | 0.041 | 0.702 | 0.036 |
| 2 | SVM | 0.727 | 0.042 | 0.739 | 0.081 | 0.657 | 0.049 | 0.720 | 0.052 | 0.647 | 0.075 | 0.678 | 0.048 |
| 3 | SVM | 0.706 | 0.056 | 0.714 | 0.061 | 0.652 | 0.050 | 0.683 | 0.081 | 0.645 | 0.055 | 0.663 | 0.067 |
| 4 | SVM | 0.728 | 0.040 | 0.726 | 0.050 | 0.654 | 0.029 | 0.703 | 0.082 | 0.653 | 0.065 | 0.671 | 0.030 |
| 5 | SVM | 0.736 | 0.045 | 0.723 | 0.061 | 0.682 | 0.016 | 0.690 | 0.056 | 0.684 | 0.039 | 0.685 | 0.034 |
| 6 | SVM | 0.723 | 0.059 | 0.716 | 0.095 | 0.654 | 0.041 | 0.700 | 0.026 | 0.649 | 0.090 | 0.670 | 0.048 |
| 7 | SVM | 0.727 | 0.023 | 0.719 | 0.056 | 0.661 | 0.054 | 0.683 | 0.048 | 0.662 | 0.031 | 0.672 | 0.037 |
| 8 | SVM | 0.739 | 0.023 | 0.749 | 0.036 | 0.664 | 0.034 | 0.720 | 0.053 | 0.656 | 0.063 | 0.683 | 0.036 |
| 9 | SVM | 0.748 | 0.046 | 0.751 | 0.083 | 0.687 | 0.076 | 0.734 | 0.067 | 0.678 | 0.105 | 0.701 | 0.079 |
| 0 | RF | 0.723 | 0.063 | 0.732 | 0.055 | 0.632 | 0.067 | 0.617 | 0.085 | 0.642 | 0.071 | 0.628 | 0.073 |
| 1 | RF | 0.717 | 0.033 | 0.704 | 0.088 | 0.666 | 0.028 | 0.657 | 0.046 | 0.676 | 0.047 | 0.664 | 0.028 |
| 2 | RF | 0.713 | 0.029 | 0.711 | 0.075 | 0.639 | 0.024 | 0.616 | 0.082 | 0.663 | 0.075 | 0.630 | 0.033 |
| 3 | RF | 0.727 | 0.048 | 0.726 | 0.041 | 0.666 | 0.058 | 0.642 | 0.095 | 0.674 | 0.069 | 0.657 | 0.083 |
| 4 | RF | 0.708 | 0.057 | 0.704 | 0.082 | 0.634 | 0.047 | 0.613 | 0.093 | 0.655 | 0.081 | 0.626 | 0.049 |
| 5 | RF | 0.730 | 0.065 | 0.718 | 0.080 | 0.673 | 0.045 | 0.655 | 0.067 | 0.685 | 0.058 | 0.668 | 0.054 |
| 6 | RF | 0.715 | 0.038 | 0.705 | 0.085 | 0.659 | 0.028 | 0.654 | 0.041 | 0.666 | 0.091 | 0.657 | 0.051 |
| 7 | RF | 0.731 | 0.028 | 0.713 | 0.064 | 0.664 | 0.045 | 0.646 | 0.035 | 0.676 | 0.050 | 0.660 | 0.039 |
| 8 | RF | 0.729 | 0.037 | 0.709 | 0.074 | 0.652 | 0.033 | 0.659 | 0.068 | 0.662 | 0.057 | 0.656 | 0.035 |
| 9 | RF | 0.739 | 0.038 | 0.731 | 0.057 | 0.664 | 0.041 | 0.639 | 0.078 | 0.679 | 0.093 | 0.654 | 0.062 |
| 0 | NN | 0.728 | 0.071 | 0.730 | 0.084 | 0.652 | 0.071 | 0.661 | 0.097 | 0.652 | 0.068 | 0.655 | 0.079 |
| 1 | NN | 0.735 | 0.033 | 0.727 | 0.059 | 0.702 | 0.028 | 0.736 | 0.050 | 0.693 | 0.038 | 0.713 | 0.034 |
| 2 | NN | 0.723 | 0.041 | 0.735 | 0.088 | 0.648 | 0.032 | 0.638 | 0.062 | 0.662 | 0.063 | 0.646 | 0.028 |
| 3 | NN | 0.701 | 0.067 | 0.711 | 0.069 | 0.643 | 0.070 | 0.646 | 0.119 | 0.641 | 0.082 | 0.642 | 0.098 |
| 4 | NN | 0.736 | 0.046 | 0.756 | 0.032 | 0.691 | 0.042 | 0.719 | 0.093 | 0.692 | 0.029 | 0.700 | 0.031 |
| 5 | NN | 0.733 | 0.045 | 0.715 | 0.059 | 0.675 | 0.024 | 0.650 | 0.055 | 0.690 | 0.038 | 0.668 | 0.036 |
| 6 | NN | 0.719 | 0.049 | 0.712 | 0.089 | 0.650 | 0.053 | 0.667 | 0.025 | 0.656 | 0.103 | 0.657 | 0.053 |
| 7 | NN | 0.726 | 0.025 | 0.717 | 0.052 | 0.666 | 0.039 | 0.699 | 0.051 | 0.660 | 0.035 | 0.678 | 0.039 |
| 8 | NN | 0.736 | 0.029 | 0.725 | 0.046 | 0.655 | 0.032 | 0.676 | 0.046 | 0.660 | 0.055 | 0.664 | 0.019 |
| 9 | NN | 0.727 | 0.035 | 0.723 | 0.080 | 0.673 | 0.029 | 0.689 | 0.029 | 0.673 | 0.074 | 0.678 | 0.040 |
